# A Unified Filtering Method for Estimating Asymmetric Orientation Distribution Functions: Where and How Asymmetry Occurs in the Brain

**DOI:** 10.1101/2022.12.18.520881

**Authors:** Charles Poirier, Maxime Descoteaux

## Abstract

Numerous filtering methods have been proposed for estimating asymmetric orientation distribution functions (ODFs) for diffusion magnetic resonance imaging (dMRI). It can be hard to make sense of all these different methods, which share similar features and result in similar outputs. The objectives of this work are two-fold: to disentangle the various filtering methods proposed in the past for estimating asymmetric ODFs, and to study the occurrence of asymmetric patterns in dMRI brain acquisitions. Hence, we describe a new filtering equation for estimating asymmetric ODFs resulting from the unification of these previously proposed filtering methods. Our method is distributed as an open-source GPU-accelerated python software to facilitate its integration into any existing dMRI processing pipeline. Following its validation on toy datasets, we apply our method to multi-shell multi-tissue fiber ODF reconstructions for 21 subjects from the Human Connectome Project in test-retest acquisitions. Our results show that our method estimates branching, fanning, bending, ending and other complex asymmetric fiber configurations in less than 2 minutes. Also, our novel number of fiber directions (NuFiD) index reveals that the filtering reduces the number of peak directions in the resulting asymmetric representation. Finally, our MNI-aligned template of asymmetries, describing the degree of asymmetry of each voxel, suggests that at least 60% of brain voxels in a dMRI acquisition contain asymmetric fiber configurations.

## 1. Introduction

In diffusion magnetic resonance imaging (dMRI), the orientation distribution function (ODF) is usually modeled as an antipodally symmetric spherical function (SF). This is motivated by the symmetric nature of the diffusion process [1]; that is, water molecules have the same probability of travelling in the direction *v* than in its opposite −*v*. The symmetric ODF is commonly expressed in a symmetric and real spherical harmonics (SH) basis [2, 3], where symmetry is enforced by including only even order SH functions in the basis and realness, by alternating between the real and imaginary part of SH functions based on its degree. However, symmetric ODFs cannot make the distinction between crossings — which account for 66-90% of white matter (WM) voxels [2, 4] — and other complex fiber configurations such as bendings, branchings and fannings [5, 6]. To address the issue, the use of asymmetric ODFs (a-ODFs) has been suggested. Indeed, by including information from neighbours into the estimation of the ODF [7, 8, 9, 10, 11, 12, 13, 14, 15, 16, 17], it has been shown that complex fiber configurations can be inferred. To represent asymmetric SF, a full SH basis [13, 14], which is obtained by adding odd-order SH functions in the basis, is often used.

There are two different categories of approaches for estimating asymmetric ODFs. On one hand, reconstruction methods [11, 13, 15, 17] attempt to fit a-ODFs onto the raw diffusion signal, by incorporating an inter-voxel regularization in the formulation of the optimization problem. On the other hand, filtering methods [7, 8, 10, 14] generate a-ODFs by applying filtering operations on an input symmetric ODF image. For all existing methods, asymmetric configurations arise as a result of including information from neighbouring voxels into the local ODF estimation. A-ODFs have been reported to depict complex fiber configurations without the need for fiber tracking [8]. These new fiber configurations are better aligned with the expected underlying anatomy [7, 13], especially in clinical settings [10, 11], where acquisitions are of low resolution. In this context, asymmetric ODFs offer great opportunities for increasing the precision of the signal without subdividing the image in smaller voxels. A-ODFs-based tractography also results in better bundle coverage and better reconstruction of bending bundles [10], increases the agreement between tractograms and the expected connectivity [13] and mitigates the infamous gyral bias [17]. Also, asymmetry spectrum imaging [15] combines a-ODFs with varying response functions to improve tractography inside the infant brain.

This article focuses on the aforementioned filtering methods. These methods can be inserted into any existing dMRI processing pipeline without the need to intervene at the ODF reconstruction step, which is typically a computationally expensive task. Also, as they do not rely on solving complex optimization problems, filtering methods are fast and can even be further accelerated via graphical processing units (GPU) programming [14]. What’s more, they can be applied on any type of spherical signal, be it diffusion ODF (dODF), fiber ODF (fODF) or any other signal expressed as a SF.

As an increasing number of different, yet similar, approaches have been suggested for estimating asymmetric ODFs, it is becoming more and more challenging to make an informed decision about the algorithm best suited to one’s particular needs. Hence, we propose a novel unified filtering equation, obtained by putting together many interesting regularizations proposed in the literature for estimating a-ODFs from any input ODF image. The proposed equation is distributed as a minimal, open-source, GPU-accelerated application. Following demonstration of the effect of our filtering equation on test datasets, we use it to study the occurrence of asymmetric configurations inside fiber ODFs reconstructed from multi-shell multi-tissue constrained spherical deconvolution [18] (MSMT-CSD). To understand *where* asymmetries occur inside dMRI brain acquisitions, we identify regions containing asymmetric ODF configurations using the asymmetry index [14, 17] (ASI), a quantity describing the degree of asymmetry of a spherical function. We then describe *how* these asymmetric configurations manifest themselves, both qualitatively through the inspection of ODF glyphs and quantitatively using our novel number of fiber directions (NuFiD) index, an extension of the number of fiber orientations [19] (NuFO) enabling classification of asymmetric ODFs. We finally generate a template describing the degree of asymmetry inside the brain tissues from test-retest acquisitions of 21 subjects from the Human Connectome Project (HCP). Our results show that asymmetric ODFs capture branching, bending, fanning and other complex asymmetric fiber configurations in 43.44% of WM voxels and 73.25% of gray matter (GM) voxels. Asymmetries are systematically identified along the gyral crowns, at the bottom of the corpus callosum (CC) and at the exit of the CC bridge.

## 2. Related works

In this section, we present previously published methods sharing a similar design for generating asymmetric ODF; the end goal being to incorporate interesting properties from these methods into a unified filtering equation. A-ODFs are obtained from a weighted sum of symmetric ODFs inside a neighbourhood following the general formulation

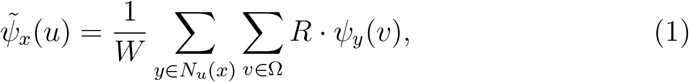

where 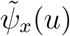 is the a-ODF at voxel position *x* for direction *u, N_u_*(*x*) is the set of the positions of the neighbours of *x*, Ω is the set of unit-length sphere directions on which the ODF is projected, *R* is the regularization weight applied to the original ODF *ψ_y_*(*v*) at voxel position *y* along direction *v* and *W* is a normalization factor. To further simplify the equations, we rewrite the vector going from current voxel *x* to voxel *y* as the multiplication of its length 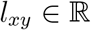 by its normalized direction 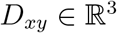, such that *y* – *x* = *l_xy_* · *D_xy_*.

### Asymmetric ODF regularization

Delputte et al [7] generate a-ODFs by regularizing a symmetric ODF field. They propose an inter-voxel regularization method which does not enforce antipodal symmetry on the ODF, resulting in asymmetries; the intuition being that an ODF at a voxel should be reflected in the ODFs of its neighbours if they belong to the same bundle. Starting from Equation 1, with the neighbourhood *N_u_*(*x*) given by the 26 immediate neighbours of *x*, the weight applied to *ψ_y_*(*v*) is

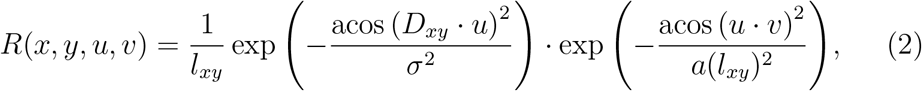

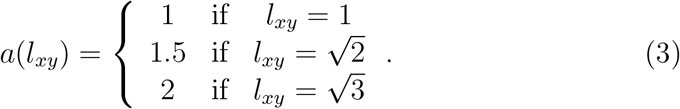

In the above equation, increasing *σ* increases the scattering of the distribution used for describing the weight assigned to each ODF value. Neighbours are weighted by their alignment to the current sphere direction *u*. The other sphere directions *v* ∈ Ω are also weighted by their alignment with the sphere direction *u*. The term *a*(*l_xy_*) flattens the bell curve as the distance between *x* and *y* increases, hence taking into account more directions for voxel *y*.

### Tractosemas

Barmpoutis et al [8] proposed *tractosemas*, a new object for describing asymmetric ODF. Obtained by diffusing ODF values across neighbours iteratively, it can be described as a weighted sum on all neighbours and directions using Equation 1. The weight *R*(*x, y, u, v*) is given by the product of 3 weights,

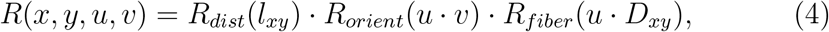

where *R_dist_* regulates by the distance between voxels, *R_orient_* by the alignment between sphere directions and *R_fiber_* by the alignment between the current sphere direction and the direction to a neighbour. The spatial weight is given by a multivariate normal distribution

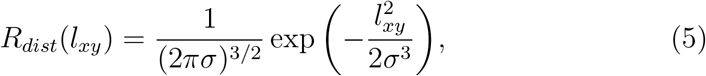

where *σ* controls the sharpness of the distribution. The orientation dependent terms are modelled as von Mises-Fisher distributions,

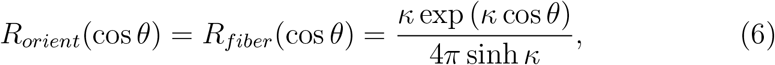

with *κ* controlling the sharpness of the distributions. In 2 to 3 iterations, the method can turn a symmetric ODF field into an asymmetric one.

### Cone-beam regularization

Asymmetric generalized cone-beam regularization (CB-REG) [10] differs from the previous methods in the way the neighbourhood is defined. The neighbourhood voxel positions are divided in two sets, *N_u_*(*x*) = *N_+u_*(*x*) ∪ *N_−u_*(*x*). The set *N_+u_*(*x*) contains all voxel positions in front of the plane defined by *u* and the second, *N_−u_*(*x*), contains the positions behind it. The neighbours positions are obtained by taking unit steps along rays launched from the center voxel *x* for each sphere directions *v* ∈ Ω. A ray is stopped as soon as the regularization weight associated to the resulting position falls below some threshold *τ* = 0.01. Because the samples are not necessarily aligned with the voxel grid, neighbours ODFs are interpolated using trilinear interpolation. Figure 1 explains the neighbourhood construction strategy visually. At the difference of the other methods presented above, the method does not average on sphere directions *v* ∈ Ω. Instead, the sphere direction considered at voxel *y* is always *D_xy_*. Hence, the filtering equation is written as

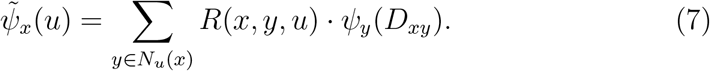

**Figure 1:**
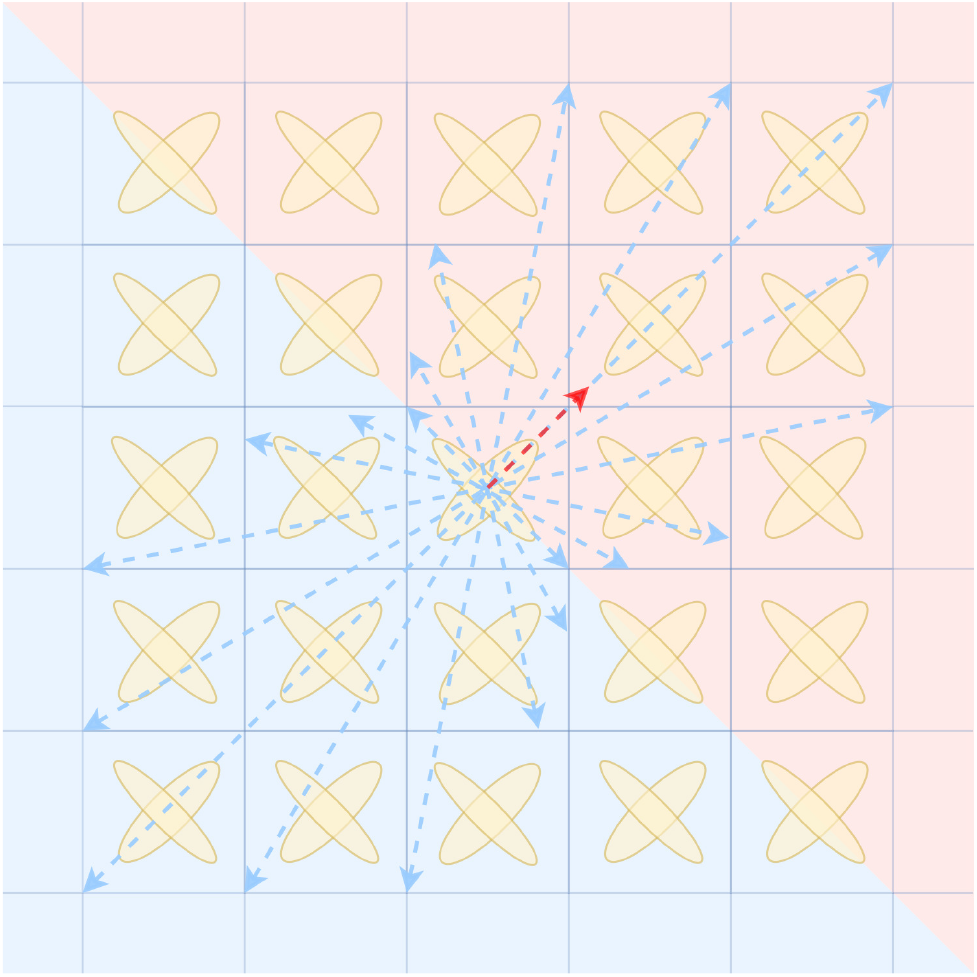
Generalized cone-beam regularization neighbourhood sampling. The current sphere direction *u* corresponds to the red arrow. The neighbourhood *N_u_*(*x*) is divided in two sets *N_+u_*(*x*) in red and *N_−u_* (*x*) in blue. Samples are drawn at each unit step along each sphere direction (blue arrows).

The regularization weight is given by

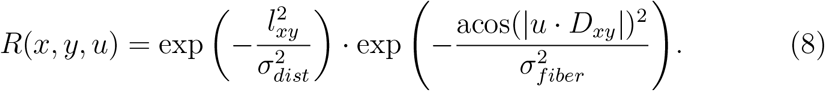

The above definition does not result in asymmetries, as it assigns the same weights to sphere directions *u* and −*u*. To get around this, the authors assign different *σ* values for neighbours depending on their belonging to *N_+u_*(*x*) or *N_−u_*(*x*). Therefore, *σ_dist_* and *σ_fiber_* are rewritten as

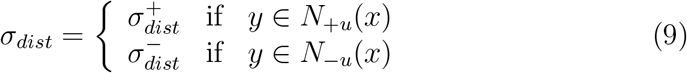

and

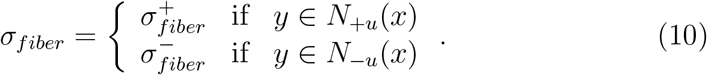

### Gaussian cone model

In [14], the neighbourhood *N_u_*(*x*) is defined as the set of all voxel positions fitting inside a cone of base radius *r* and height *h* aligned with *u* (see Figure 2). Only the sphere direction *u* is used in the averaging, removing the sum on *v* ∈ Ω from Equation 1. The Gaussian cone model assigns a weight to neighbours based on their distance to the center axis of the cone. Starting from the definition of a zero-centered Gaussian distribution with standard deviation *σ*,

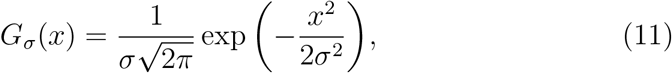

the regularization for the Gaussian cone model is given by

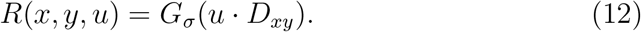

**Figure 2:**
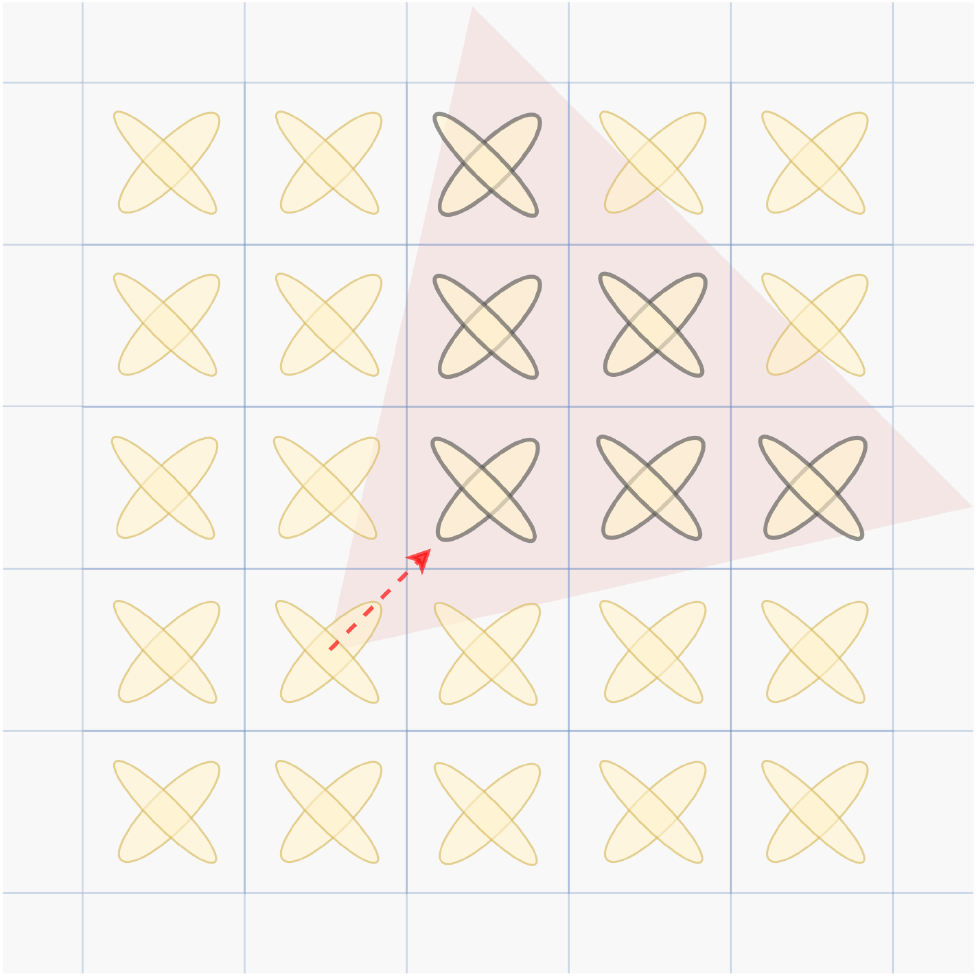
Cone-based neighbourhood used in [14]. The cone corresponding to some direction *u* is represented by the red triangle and only the ODFs covered by the cone (black outline) are included in the neighbourhood *N_u_*(*x*).

This filter only weights neighbours by their alignment with the direction *u*, not taking into account the distance between *x* and *y*.

### Bilateral cone filter

The bilateral cone filter [14] is inspired by the popular edge-preserving bilateral filter [20] for 2D image processing. Using the same definition for *N_u_*(*x*) as in the Gaussian cone model, its extension to spatio-angular images is given by

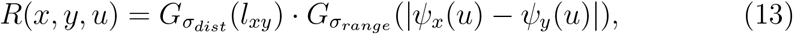

where *G_σ_* follows the definition in Equation 11. The first distribution, *G_σ_dist__*, weights neighbours by the distance between *x* and *y*, while the second, *G_σ_range__*, weights neighbours by the difference between intensities *ψ_y_*(*u*) and *ψ_x_*(*u*). This later regularization has the effect of preserving edges by giving lesser weights to values dissimilar to the current ODF value.

## 3. Methods

From the methods presented above, we design a novel unified filtering equation for estimating asymmetric ODFs, which we later use to investigate the occurrence of asymmetric patterns inside diffusion-weighted brain acquisitions. This section describes the methods used to achieve our objectives.

### 3.1. A unified filtering equation for a-ODFs

We propose a new unified filtering method for estimating asymmetric ODFs, inspired by the methods presented in the previous section. In the works described above, we saw that neighbours are weighted according to:

1. The distance between the processed voxel *x* and its neighbour *y*, *R_spatial_*.
2. The distance between the direction *D_xy_* from the voxel *x* to its neighbour *y* and the currently processed unit direction *u*, *R_align_*.
3. The distance between the processed direction *u* and the currently considered direction *v* on the input ODF image, *R_angle_*.
4. The difference between the intensities of the ODFs *ψ_x_*(*u*) and *ψ_y_* (*v*), *R_range_*.

We encompass these regularization weights into a single asymmetric filtering equation, in the form

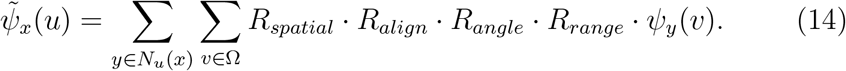

The filtering methods presented in section 2 can still be expressed by adapting this new equation. Table 1 summarizes how each method fits in our new equation. Our method is the only one to support all four regularizers at the same time.

**Table 1:** Comparison of the main features of each described filtering method.

| Method | $N_u(x)$ | $R_{spatial}$ | $R_{align}$ | $R_{angle}$ | $R_{range}$ |
| --- | --- | --- | --- | --- | --- |
| ODF postprocessing [7] | $3 \times 3 \times 3$ window (excluding self) | None | Gaussian | Gaussian | None |
| Tractosemas [8] | Window, centered on $x$ | Gaussian | von Mises-Fisher | von Mises-Fisher | None |
| CB-REG [10] | Ray marching along each $v \in \Omega$ until $R(y, v) < \tau$ | Gaussian | Gaussian (considered direction $v$ at $y$ is $D_{xy}$) | | None |
| Gaussian cone filter [14] | Voxel inside cone aligned on $u$ | None | Gaussian | None | None |
| Bilateral cone filter [14] |  | Gaussian | None | None | Gaussian |
| Our unified filter | Window of half-width $\lfloor 3 \cdot \sigma_{spatial} + 0.5 \rfloor$ | Gaussian | Gaussian | Gaussian | Gaussian |

Because they are widely used in 2D image processing, we model each weight using a normal distribution with standard deviation *σ*. The resulting filtered ODF is then given by

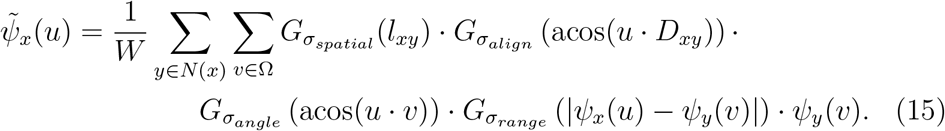

It is worth noting that, because it is related to the image intensities, the *σ_range_* parameter should be set relative to the range of amplitudes, *ψ_range_* = *ψ_max_* – *ψ_min_*, in the processed image. The neighbourhood *N*(*x*) is a window centered around *x* with a half-width of 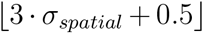 to include all voxels within 3 standard deviations of the current voxel. The voxel *x* is included in *N*(*x*). Because *R_align_* depends on the direction to neighbours, we get *D_xy_* = *D_xx_* = 0 when computing the weight for *x*, resulting in *u* · *D_xx_* = 0 and 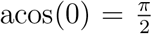. As the voxel *x* is not actually at a distance of 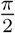 from *u*, we enforce that the voxel *x* is perfectly aligned with *u* when building the alignment filter. Individual filters can be disabled. In the cases of *R_spatial_*, *R_align_* and *R_range_*, it is equivalent to setting them to 1, while in the case of *R_angle_*, we have

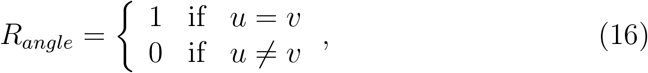

which is equivalent to removing the sum on *v* ∈ Ω and setting *v* = *u*. Our method is designed this way so that applying our filter with all four regularizers disabled is equivalent to applying a mean filter on independent sphere directions, which is how we would filter a multi-channel color image in classical image processing [21].

### 3.2. Quantifying the asymmetry of a-ODFs

To measure the degree of asymmetry of the filtered a-ODFs, we use the measure from [14], which is based on the cosine similarity between the ODF *ψ_x_*(*u*) and its flipped version *ψ_x_*(−*u*). Intuitively, if the ODF is perfectly symmetric, then *ψ_x_*(*u*) = *ψ_x_*(−*u*) results in a cosine similarity of 1. The cosine similarity is given by

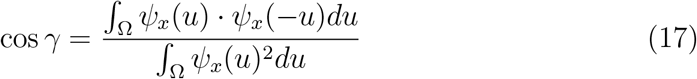

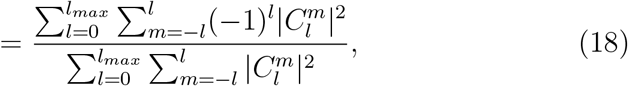

where 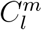 is the SH coefficient of order *l* and degree *m* describing the ODF. Then, from cos *γ*, we can compute the asymmetry measure *α* using

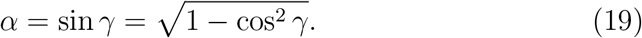

From now on, we will refer to this quantity as the *asymmetry index* (ASI), as was suggested in [17].

### 3.3. From NuFO to NuFiD

When working with symmetric ODFs, the number of fiber orientations [19] (NuFO) is often used to summarize fiber configurations at the voxel level. This is done by counting the number of peaks extracted from the ODF, based on some absolute and relative thresholds, and on a minimum separation angle. The absolute threshold is the minimum amplitude a peak direction must have to be valid and the relative threshold is a fraction of the range of amplitudes of the processed ODF a peak direction must have to be valid. In the case of symmetric ODF, it is enough to consider opposite directions *v* and −*v* as a single, coherent orientation and to extract peaks on a single hemisphere. However, this assumption does not hold when working with asymmetric SF, as a given orientation can have two distinct values based on the hemisphere from which it is sampled. Hence, we extend the NuFO for the case of a-ODFs by proposing a measure of the number of fiber directions (NuFiD). This novel NuFiD index is obtained by counting the number of maxima on a whole sphere instead of a single hemisphere. Therefore, a symmetric fODF with a given NuFO has a NuFiD index of twice this value. This implies that a perfectly symmetric fODF is restricted to having an even NuFiD index. Possible fiber configurations for NuFiD of 1 to 6 are shown in Figure 3.

**Figure 3:**
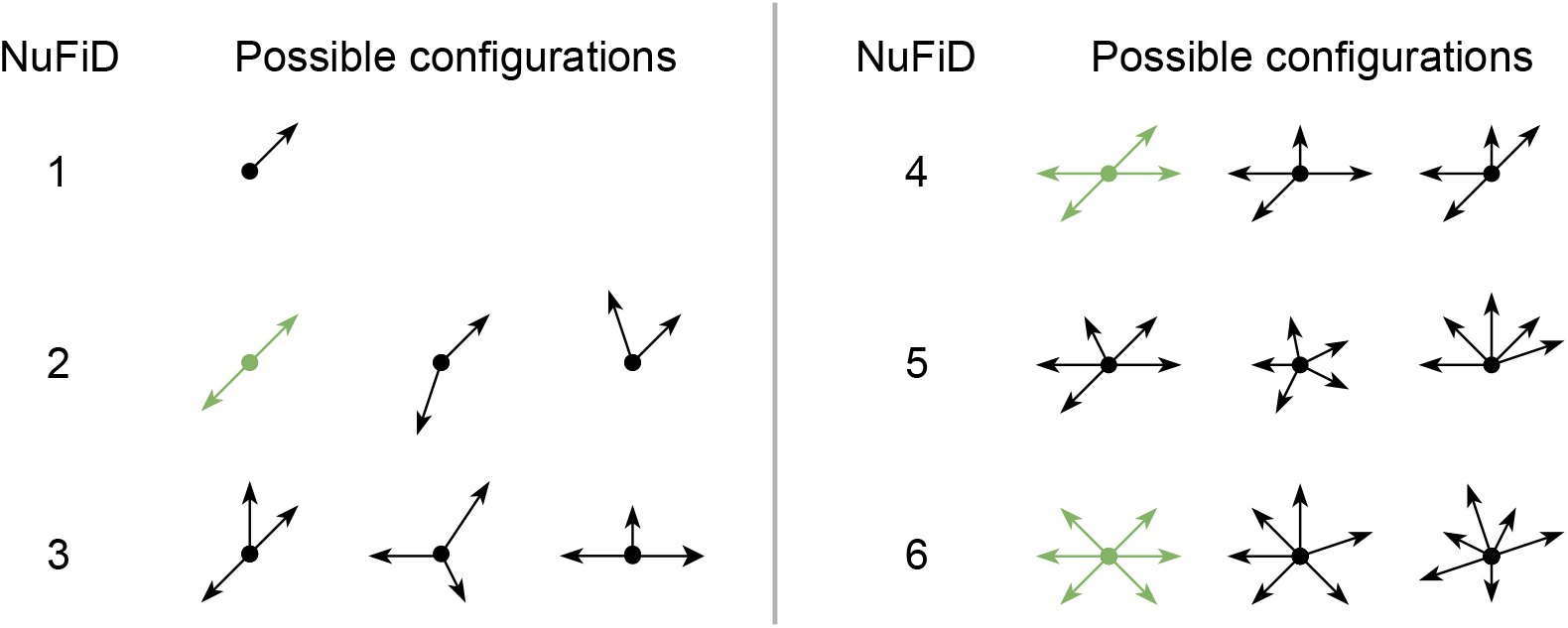
Fiber configurations for NuFiD indices of 1 to 6. Symmetric configurations for NuFO of 1, 2, 3 (NuFiD of 2, 4, 6, respectively) are shown in green.

### 3.4. Implementation details

Our method is implemented in python and accelerated using OpenCL for python [22] and Numba [23]. We deal with edges by zero-padding the input SH image. The application automatically detects the maximum order of the SH basis in which the data is given. To meet the high memory requirements needed for storing SF coefficients for a whole brain, SH coefficients images are processed in batches — 40 × 40 × 40 sub-images. SH patches are converted to SF amplitudes by projecting them on 200 evenly-distributed directions on the unit sphere. Each SF sub-image is then copied on the GPU, where each processing unit is responsible for computing the filtered SF amplitudes inside one voxel. The output SF amplitudes are then copied back to the CPU where they are converted to SH coefficients using a full SH basis with same maximum order as the input. Our application is distributed as a standalone, minimal, open-source python package at https://github.com/CHrlS98/aodf-toolkit.

### 3.5. Experiments

To better understand the behaviour of each regularizer contributing to the filtering, our filter is first applied to test datasets. Then, the method is applied to real data to investigate where and how asymmetries occur inside brain tissues. In this section, a detailed explanation of the experiments carried out in this work is given.

#### 3.5.1. Understanding the role of each filter

We first evaluate the individual effect of each regularization term implicated in Equation 15 qualitatively. Each regularizer is applied on its own after disabling the three other regularizers. The first dataset used for the experiment is a fODF image (maximum SH order 8) reconstructed by nonnegativity constrained spherical deconvolution [24] using the b-1500 shell of the averaged DWI acquisitions for the Fibercup phantom [25, 26, 27, 28]. We simulate a second test dataset consisting of fully isotropic ODFs of varying amplitude (0.0, 0.56, 1.41) distributed on a 5 × 5 × 5 grid. Only the middle slice of the simulated data contains non-null ODFs. For baseline comparison, a mean-filtered version of the datasets inside a 3 × 3 × 3 window is also computed by disabling all four regularization terms. At the exception of the experiments regarding spatial regularization where the window width depends on the choice for *σ_spatial_*, the filtering window width is set to 3 × 3 × 3 across all experiments.

#### 3.5.2. Asymmetric patterns in MSMT-CSD a-fODFs

MSMT-CSD fiber ODFs [18] of maximum SH order 8 are reconstructed for 21 subjects from the Human Connectome Project [29] (HCP) in test-retest (42 acquisitions in total) using DIPY [30]. We used all the HCP preprocessed datasets [31] and used Tractoflow [32] processes for denoising, N4 correction, and brain extraction. Asymmetric fODFs (a-fODFs) are then estimated for each subject using *σ_spatial_* = 1.0, *σ_align_* = 0.8, *σ_angle_* = None, and *σ_range_* = 0.2 · *ψ_range_*. Filtering parameters are chosen thanks to the experiments from section 3.5.1 and validated qualitatively on a subset of our subjects. Voxels where there is no signal in the input fODF image are set to 0 in the output a-fODF image. ASI maps are computed for each resulting image. Symmetric and asymmetric fODF glyphs are also studied in the `dmri-explorer` application [33]. NuFiD maps are generated using an absolute threshold of 0.1, a relative threshold of 0.3 and a minimum separation angle of 25 degrees for both the input fODF images and the filtered a-fODF images. The thresholds are chosen experimentally to obtain a NuFiD index of 2 (NuFO of 1) in the corpus callosum (CC) and a NuFiD index of 6 (NuFO of 3) in the centrum semiovale on the symmetric fODF images. There are therefore two NuFiD maps per acquisition: one computed from the input symmetric fODF image and another one computed from the a-fODF image resulting of the filtering. For each pair of NuFiD maps, we compute the distribution of a-fODF NuFiD indices inside some masks. These masks are obtained by taking all voxels with a given NuFiD index of *m* prior to filtering (in the input fODF NuFiD map), for *m* ∈ {0, 2, 4, …, *m_max_*} with *m_max_* the highest value in the fODF NuFiD map. A visual explanation of this experiment is given in Figure 4. Finally, average NuFiD distributions and their standard deviations (std) are computed.

**Figure 4:**
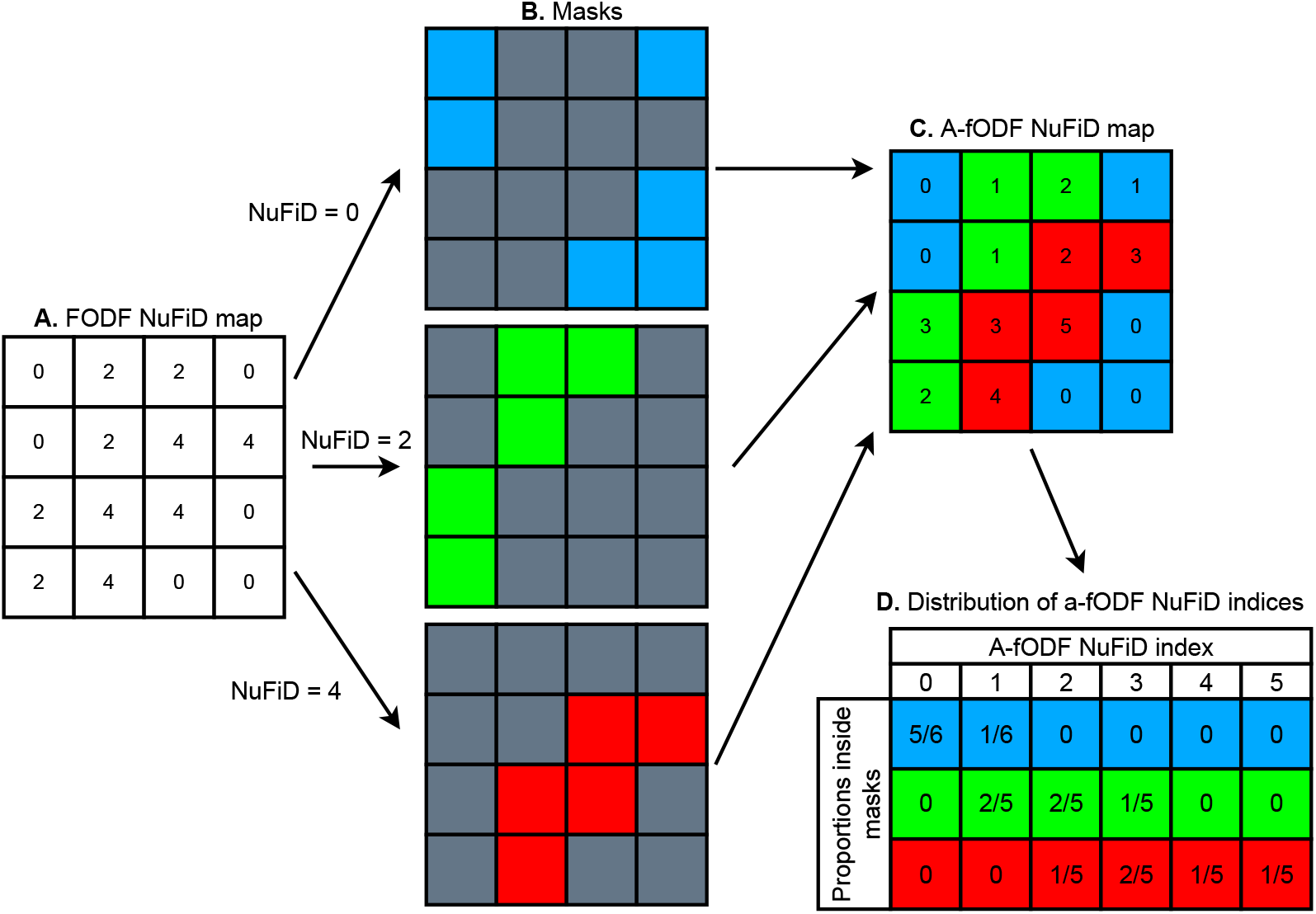
Computing the distributions of NuFiD indices. (A.) NuFiD map computed from the symmetric fODF image for a given acquisition. (B.) Masks corresponding to each NuFiD value found in the input NuFiD map. (C.) A-fODF NuFiD map for the same acquisition. (D.) Distribution of a-fODF NuFiD indices inside each mask.

#### 3.5.3. Template and proportion of asymmetries

A template describing the ASI of brain voxels is also created. As part of the HCP preprocessing pipeline, the diffusion-weighted images are registered to 1.25 mm^3^ resampled T1-weighted images. These resampled T1-weighted images are registered to the ICBM 2009a non-linear asymmetric template from the Montreal Neurological Institute (MNI space) [34, 35]. Resulting linear and non-linear transformations are then applied to the asymmetry maps using trilinear interpolation and the resulting registered images are averaged together. The standard deviation of the asymmetry degree per voxel is also computed. The resulting image is masked using a brain mask from the MNI template. Brain voxels are classified using the MNI tissue probability maps by assigning to each voxel its tissue of highest probability. Then, WM and GM masks are generated by taking all voxels belonging to the WM and GM class, respectively. A mask containing all WM and GM voxels is also obtained from the union of both masks. Inside each mask, the proportions of voxels with an ASI higher than some threshold *τ* are reported for *τ* ∈ [0.0, 1.0] by steps of 0.05. Finally, the proportions of asymmetric voxels inside each mask are reported by thresholding the ASI template to 0.35 using MI-Brain [36].

## 4. Results

In this section, we describe the results of each experiment. In subsection 4.1, the effects of the filter weights are described qualitatively. Asymmetric patterns found inside MSMT-CSD a-fODFs following filtering are reported in subsection 4.2. The resulting template of asymmetry is studied in subsection 4.3.

### 4.1. The effect of filtering

In Figure 5, the effect of spatial filtering for *σ_spatial_* ∈ {0.6, 1.0} (with respective window widths of 5 and 7) is compared to the input and mean-filtered images for a region of interest showing three bundles splitting. Only voxels inside a WM mask are shown. To concentrate on the shape of the SF, each ODF is normalized by its maximum amplitude. As *σ_spatial_* increases, we see that the ODFs lobes are progressively better aligned with those of their neighbours, enhancing inter-voxel coherence at the cost of losing spatial resolution. Also note that for *σ_spatial_* = 1.0, the output SF are very similar to the mean-filtered ones.

**Figure 5:**
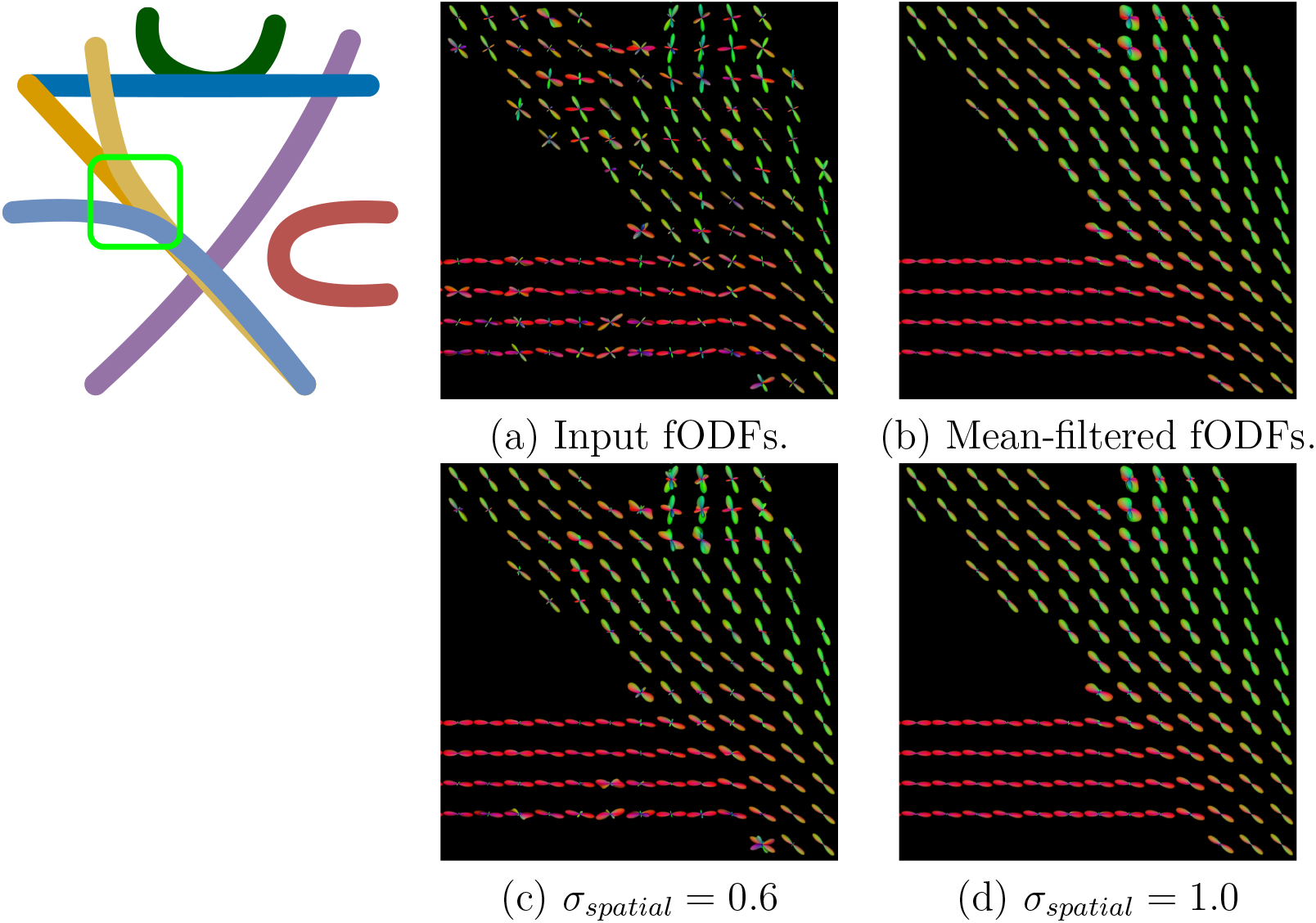
The effect of spatial regularization for varying *σ_spatial_* and varying window size for the regions identified by the green box on the left.

In Figure 6, the effect of angle filtering is shown on the Fibercup phantom inside a 90 degrees crossing for *σ_angle_* ∈ {0.2, 0.4, 0.6}. Angle filtering averages neighbouring sphere directions between themselves. As a consequence, increasing *σ_angle_* results in a loss of angular resolution as ODF amplitudes get blurred together. Also, we see that even the slightest averaging between sphere directions has a huge impact on the angular resolution of ODF and does not result in better inter-voxel coherence compared to a simple mean-filtering.

**Figure 6:**
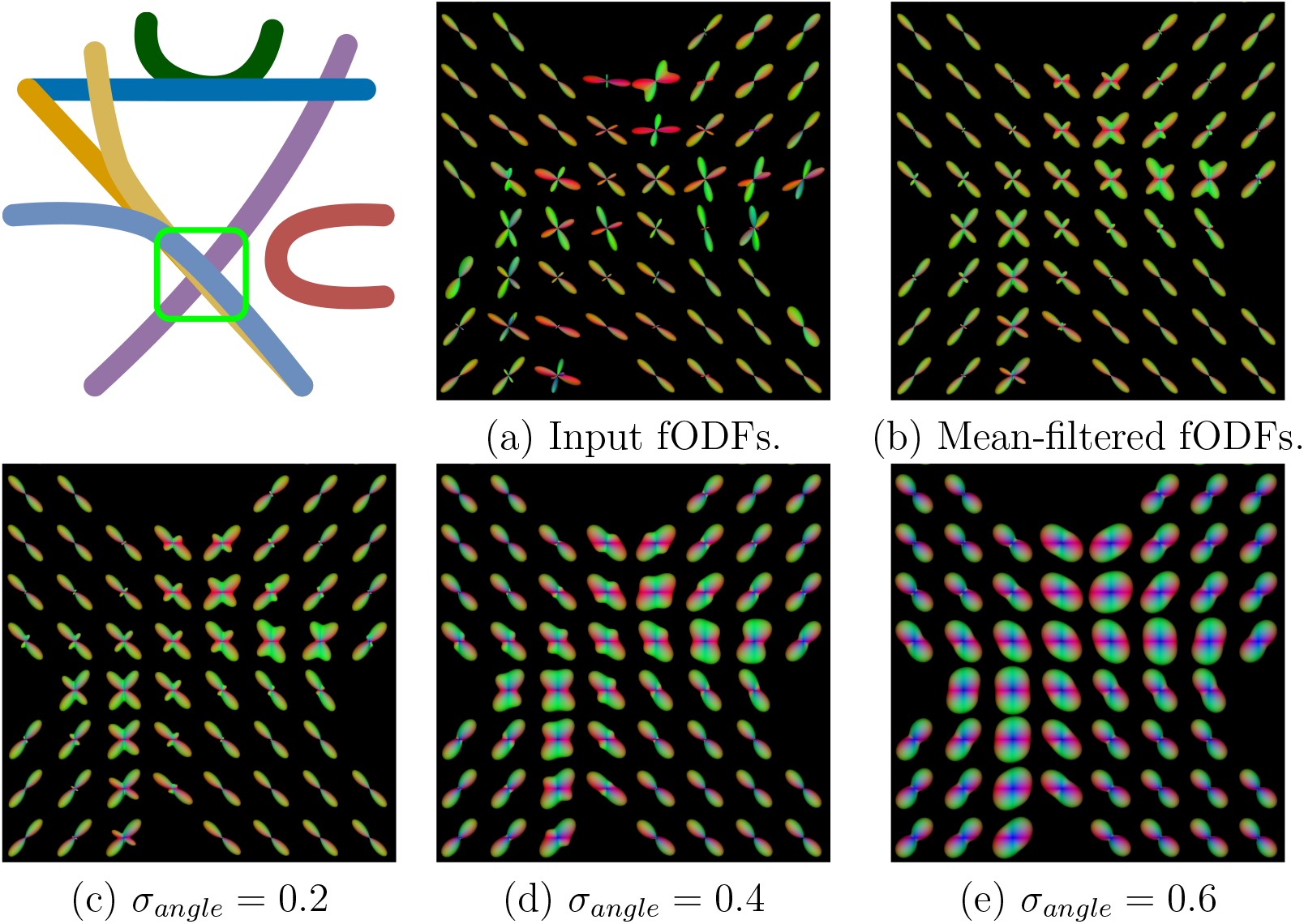
The effect of angle regularization for varying *σ_angle_* inside a 3 × 3 × 3 window. The region of interest is identified by the green box on the drawing in the top left corner.

The edge-preserving properties of the range filter are demonstrated in Figure 7 for *σ_range_* ∈ {0.1, 0.2, 0.4} · *ψ_range_* on the isotropic ODFs dataset. As *σ_range_* increases, we see that voxels containing ODFs highly dissimilar from their neighbours are more and more affected by the averaging. On the contrary, when *σ_range_* is set to a very small value, the weights assigned to any neighbour are too low to have an impact on the output image. This is observed for *σ_range_* = 0.1 · *ψ_range_*. Hence, one must be careful when selecting a value for *σ_range_*, as a too low value may cancel the effect of the other regularizers applied on the image.

**Figure 7:**
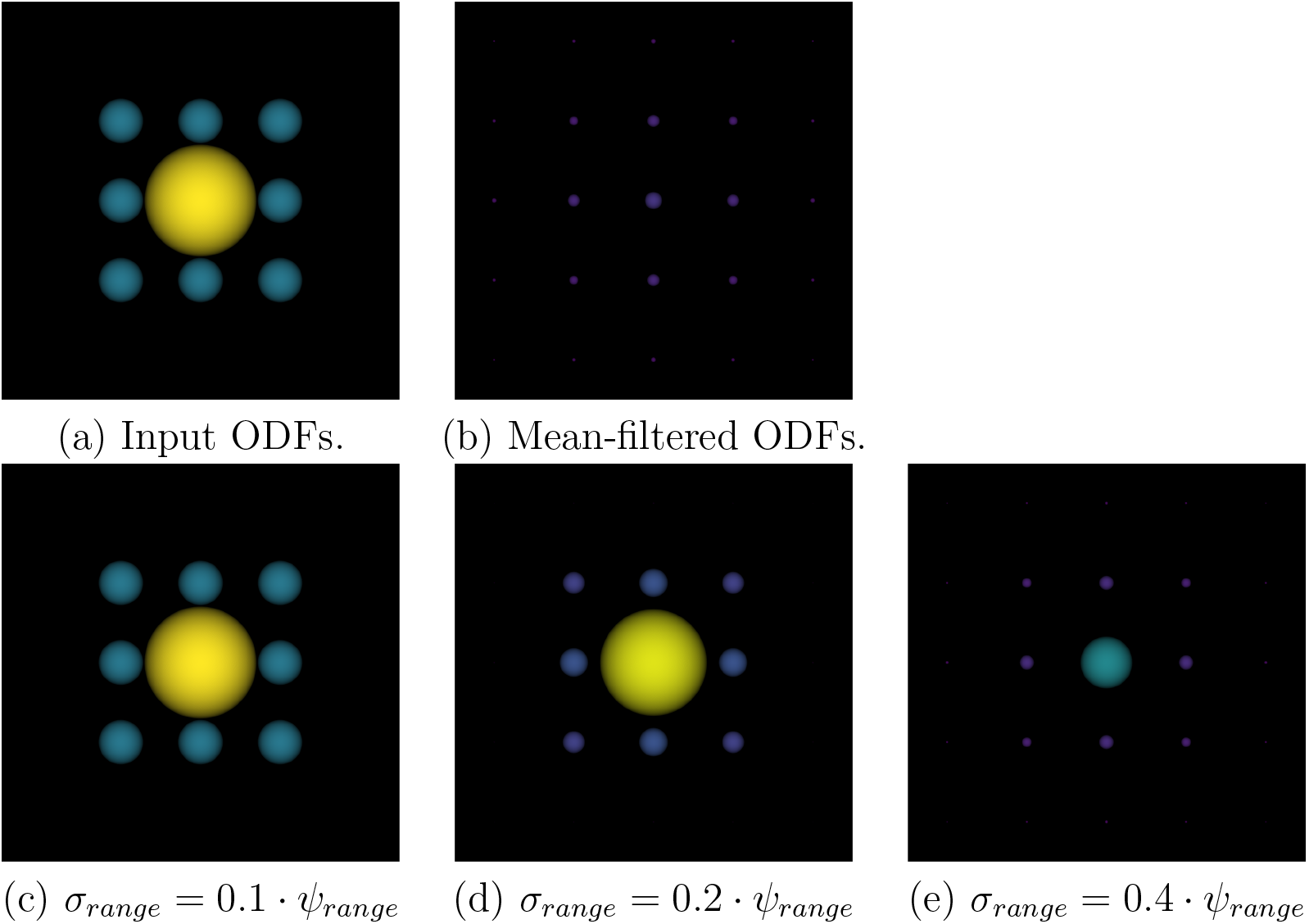
Range filtering applied on the simulated isotropic ODFs dataset. In the input image, the center ODF has an amplitude of 1.41 and its neighbours have an amplitude of 0.56. The top row shows the input and mean-filtered images while the bottom row shows the result of filtering for various values of *σ_range_*.

At the difference of all the previous regularization weights, alignment filtering generates asymmetric SF. This behaviour is shown in Figure 8 on the C-shaped bundle of the Fibercup phantom for values of *σ_align_* ∈ {0.2, 0.6, 1.0}. Regularizing on the alignment between the current direction and the direction to a neighbour results in different filters for opposite directions *u* and −*u*. Indeed, as the angle between *u* and *D_xy_* increases, the resulting weight applied at voxel *y* decreases. We see a behaviour similar to spatial regularization where an increasing value for *σ_align_* results in progressively smoother transitions between voxels; more neighbours have a contribution of significance to the filtered ODF. Furthermore, we see bending asymmetric ODFs for all tested parameter value.

**Figure 8:**
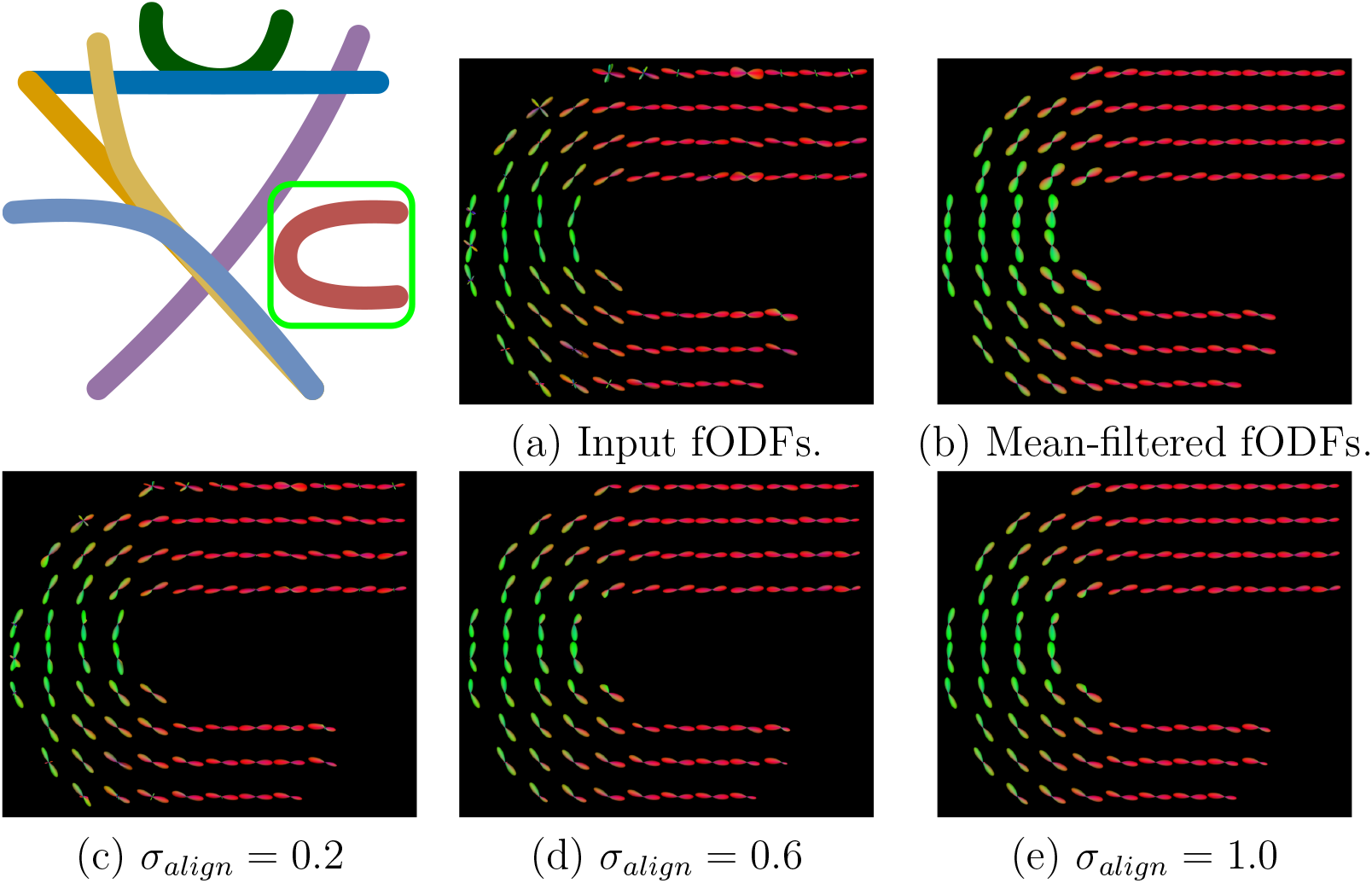
The effect of alignment regularization for varying *σ_align_* using a 3 × 3 × 3 window. The region of interest in identified by the green box in the top left drawing.

### 4.2. Asymmetric patterns in MSMT-CSD a-fODFs

As seen in Figure 9, our filtering equation successfully turns a symmetric fODF field from real in-vivo data into an asymmetric one. The figure shows the asymmetry map for a subject as well as symmetric and asymmetric glyphs for four regions identified by colored boxes. We also report the ASI for various a-fODF configurations found in the data. We see that the ASI is highly sensible to asymmetric configurations. Indeed, the bottom-left glyph, displaying very little asymmetries, has a reported ASI of 0.17. However, looking closely, we see some differences between the two smallest fODF lobes. Also, the other glyphs, displaying more obvious asymmetric configurations, have an ASI of 0.40 and above.

**Figure 9:**
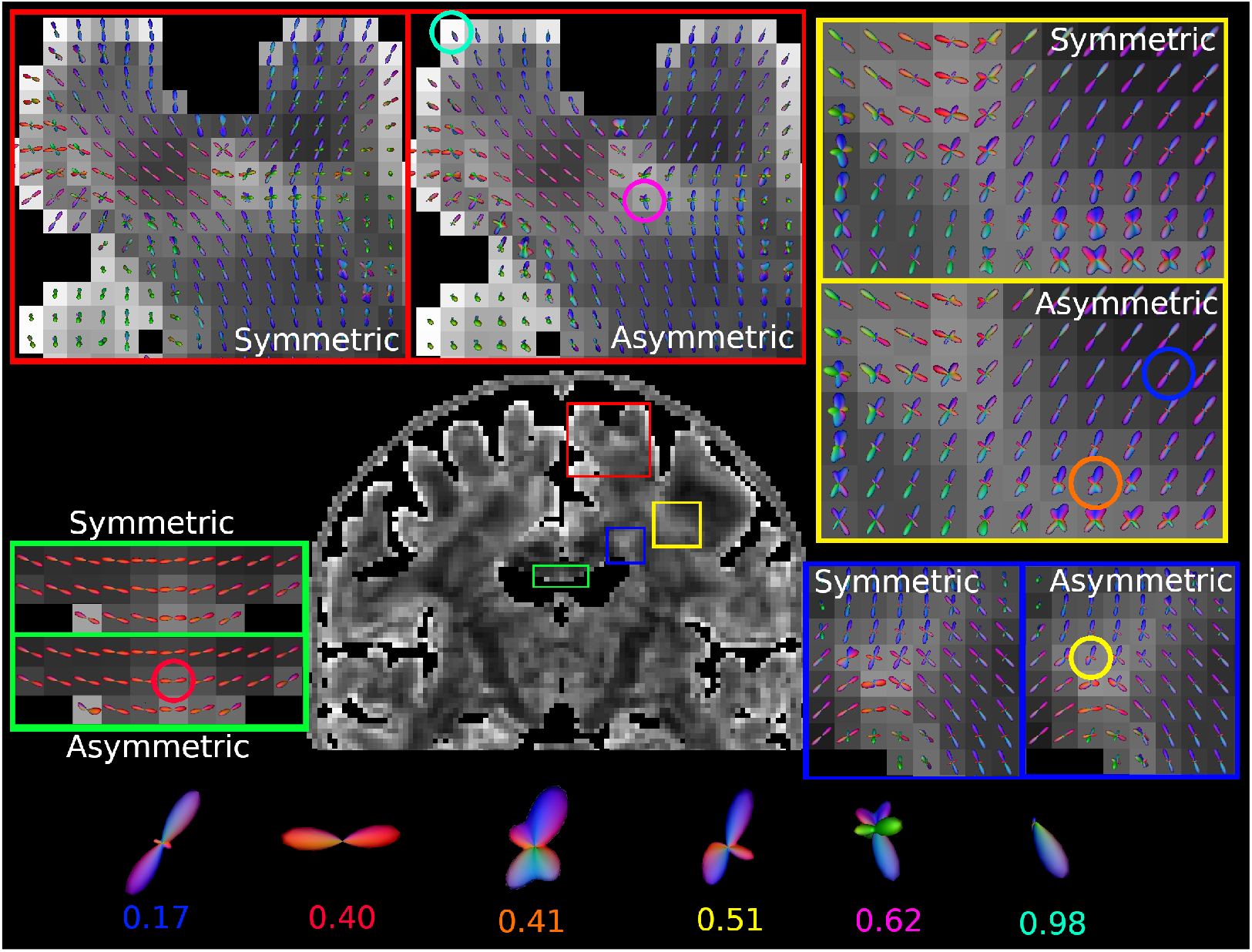
Comparison of fODF configurations before/after filtering for a single subject `(111312_re)`. The glyphs are overlayed on the asymmetry map. Interesting glyphs identified by a colored circle are shown on the bottom row with their corresponding ASI.

The red window shows a zoomed-in gyrus in which we see asymmetric fODFs capturing branching and bending fiber trajectories. We also see that single-fiber a-fODFs close to the cortex have a lobe of smaller amplitude in the direction facing gray matter than in the direction going inward. Hence, these endpoint a-fODFs capture the ending of fiber tracks as they reach the cortex. Also note how the asymmetry degree is at its highest along the cortex. This is a consequence of the highly asymmetric neighbourhood in this region, consisting of brain tissue on one side and cerebrospinal fluid (CSF) on the other.

We also report slightly bending a-fODFs in the bottom-most portion of the CC, as can be appreciated in the green box. These bent configurations better capture the U-shaped nature of fibers in the bridge of the CC as well as the repulsion of fiber trajectories from the CSF voxels.

Asymmetric configurations also appear deeper inside the WM. The yellow frame shows asymmetric glyphs in a region of crossing fibers. Complex asymmetric fiber configurations arise from the filtering as neighbourhood information is taken into account at the voxel level. This gives interesting results in voxels originally containing fODFs with low angular details (see the circle in orange), for which the filtering has a sharpening effect. Furthermore, the filtering modifies differently the amplitudes of antipodal sphere directions in crossing fODFs, resulting in a-fODFs tending toward branching configurations.

The blue frame shows a region at the exit of the CC. We notice the emergence of branching and fanning fiber configurations in voxels initially containing one or two principal fiber directions. Because some directions in the symmetric fODF are not supported by the neighbourhood, the filtering removes the misleading fODF lobes from the estimated a-fODF (see the circle in yellow), simplifying the interpretation of the a-fODF configurations for this region.

Figure 10 shows the ASI maps for three subjects from our cohort. We see that there is an inter-subject agreement in the ASI of brain regions. Indeed, the cortex “lights up” in red for all three displayed images, the bottom-most section of the CC displays asymmetries across all subjects, the exit of the CC is consistently reported as asymmetric and similar contrasts are observed inside the WM for all 3 subjects as well.

**Figure 10:**
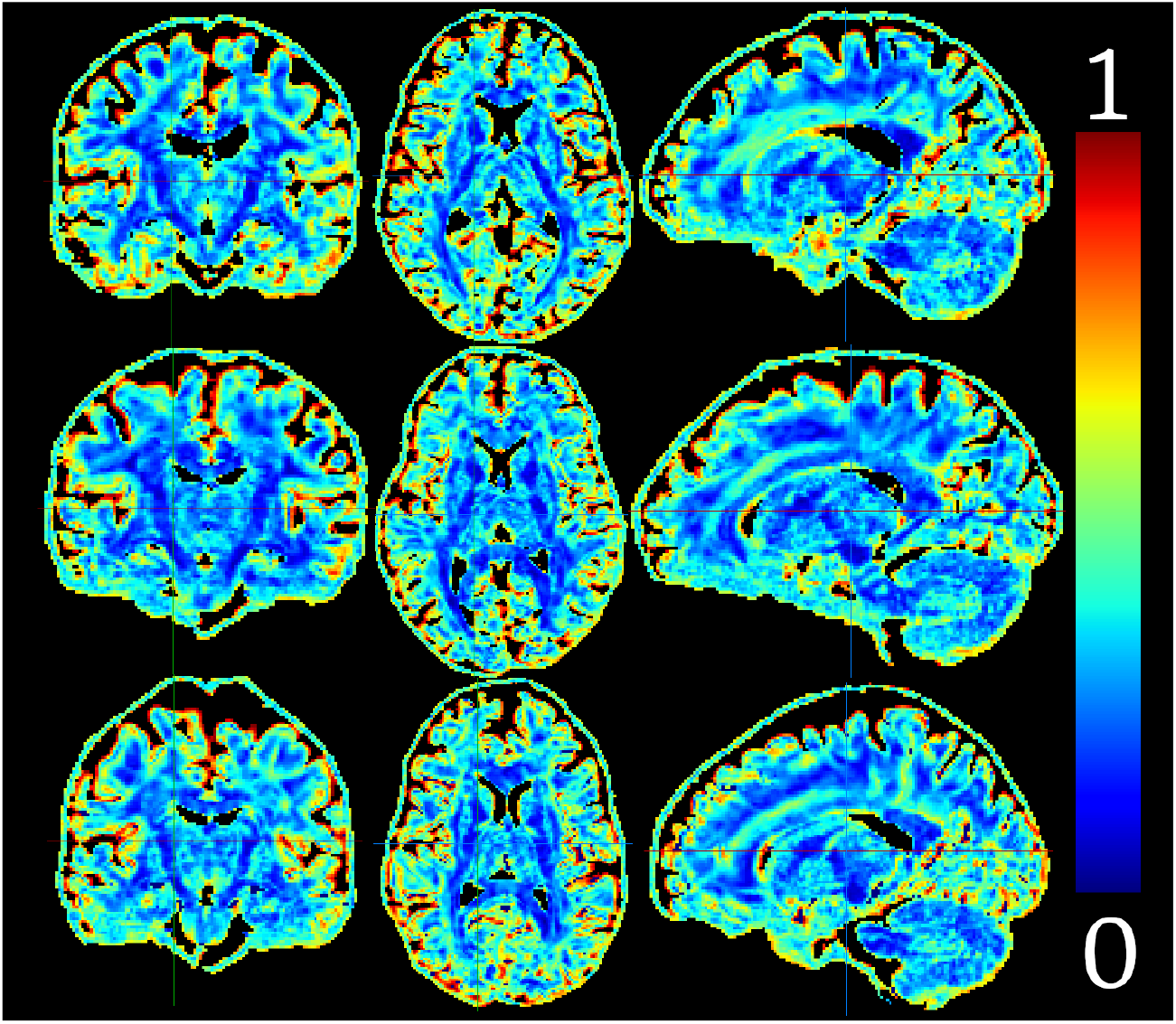
Asymmetry maps for three subjects (top: `111312_re`, middle: `130518_re` and bottom: `200614_re`). Asymmetric regions are consistently observed across subjects.

We also show the NuFiD maps for one subject in Figure 11, computed from the input fODF on the left and from the a-fODF image on the right. The NuFiD map extracted from the symmetric fODF image contains only even order NuFiD values, as each maxima found on an hemisphere also exists on the opposite hemisphere. We see the emergence of odd NuFiD values post-filtering, meaning that the resulting a-fODFs have different values for opposite directions. The cortex, highly asymmetric, has a NuFiD value of 1, consistent with the endpoint a-fODFs observed in Figure 9. We also see that odd NuFiD are observed at the interface of regions of different NuFiD value — values of 1 appear between values of 0 and 2, values of 3 appear between 2 and 4, and so on. In particular, we report 3-NuFiD a-fODFs at the exit of the CC, where we previously reported asymmetric fiber branchings. Lastly, we report an overall decrease in the NuFiD of a-fODFs compared to fODFs. Indeed, there are less 6- and 8-valued voxels following the filtering, suggesting that removing the symmetry constraint successfully removes misleading fiber directions which are not supported by the neighbourhood.

**Figure 11:**
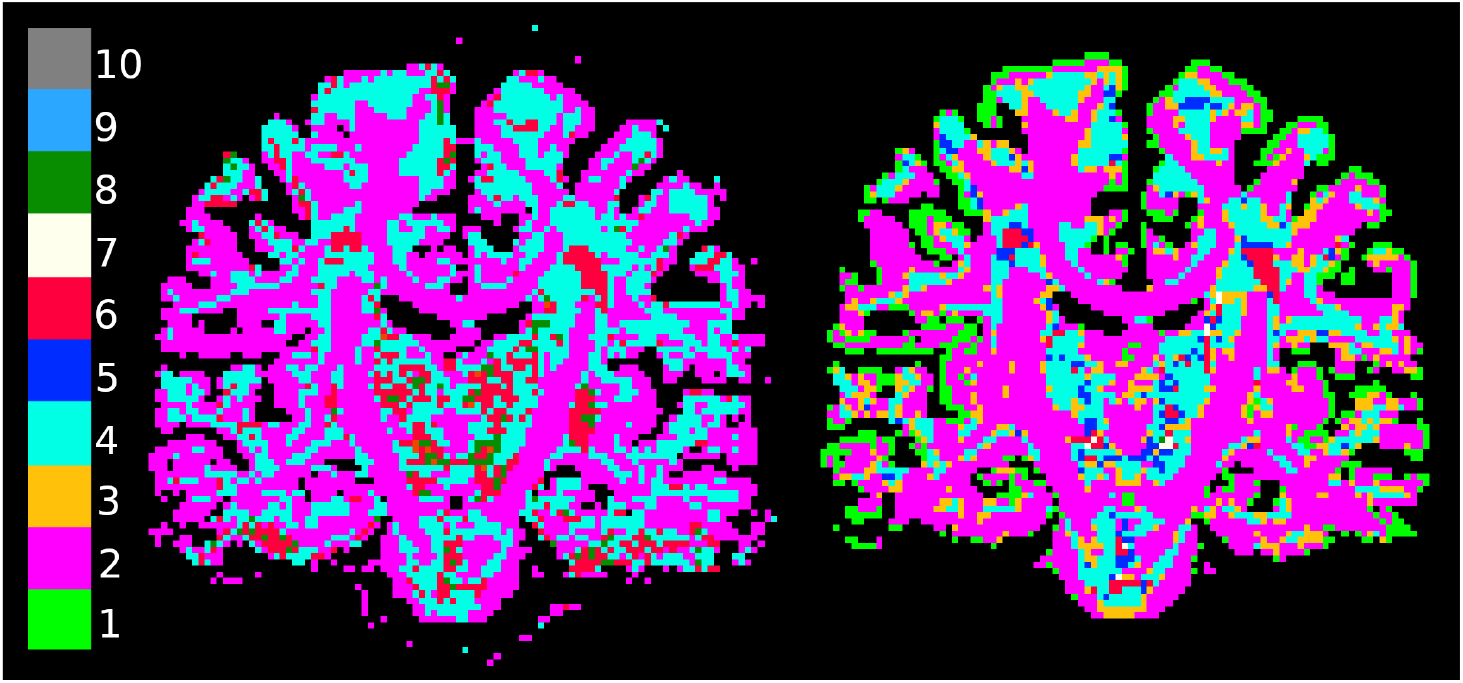
NuFiD maps computed from fODF images (left) and a-fODF images (right) for subject `200614_re`. Recall that for even NuFiD indices, `NuFO = NuFiD/2`.

The distributions of NuFiD following the filtering are reported in Figure 12 separately for test (top row) and retest (bottom row) acquisitions. Each curve shows the distribution of a-fODF NuFiD indices inside the set of voxels with a NuFiD index of *m* prior to the filtering (in the symmetric fODF image). The curves are drawn for *m* ∈ {0, 2, 4, 6, 8, 10}. The standard deviations, reported as error bars, show that the reported proportions are consistent across subjects. Our results in test and retest show that the experiment is also reproducible across acquisitions. These results show that the effect of the filtering on the distribution of NuFiD indices is consistent across subjects and acquisitions.

**Figure 12:**
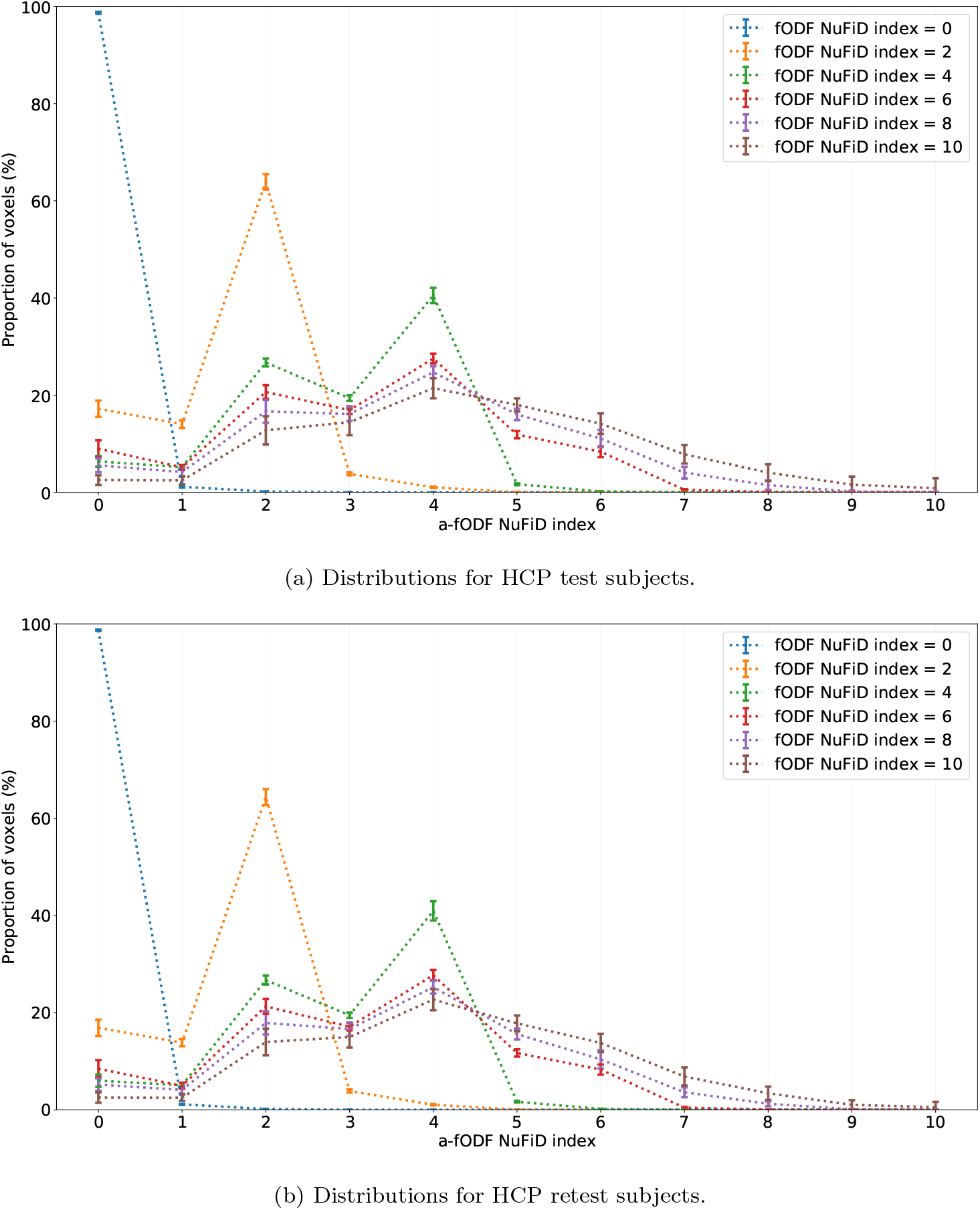
A-fODF NuFiD mean distributions and standard deviations inside voxels with a given symmetric fODF NuFiD index. Each curve describes the proportion of voxels among those having a NuFiD index of *m* prior to the filtering (*y* axis) ending up with a given NuFiD index (*x* axis) posterior to the filtering.

The blue curve shows the distribution of a-fODF NuFiD indices inside voxels with a NuFiD index of 0 in the input fODF image. Almost all of the voxels keep their value following the filtering, a direct result of the output a-fODFs being clipped to 0 in voxels where the input image is null. A small proportion of these 0-valued input voxels have a NuFiD index of 1 after the filtering, characteristic of endpoint a-fODFs found in voxels bordering CSF voxels.

The orange curve shows the distribution of a-fODF NuFiD indices for voxels with a fODF NuFiD index of 2. These corresponds to voxels with a NuFO of 1, corresponding to a single-fiber population. Once again, a majority (around 60%) of the filtered a-fODFs keep their initial NuFiD index. We also report that for almost 20% of these voxels, the NuFiD index in the a-fODF image ends up being null. In another approximate 15% of cases, the NuFiD index get demoted to 1. This is consistent with the behaviour observed along the cortex on the single subject shown in Figure 11, where NuFiD indices of 1 or 0 replace NuFiD indices of 2 due to the 0-valued fODFs found in the CSF. Because the amplitudes of fODFs get progressively smaller as we reach deeper into the gray matter, averaging these fODFs with CSF voxels attenuates one or both fODF peaks enough to result in a change of NuFiD value.

From the green curve, we see that approximately 40% of voxels with a fODF NuFiD index of 4 keep this value after the filtering. As we recall, a symmetric fODF with a NuFiD index of 4 corresponds to a crossing of 2 fiber populations (NuFO = 2). In many cases, however, we see that taking into account information from neighbours changes the amplitudes of the resulting a-fODF enough to remove 1 or 2 principal fiber directions. Therefore, following the filtering, there are more voxels that change value than there are that keep the same NuFiD index.

In red, we see the distribution of a-fODF NuFiD indices inside voxels with a NuFiD index of 6 in the symmetric fODF image. There is an important decrease in the proportion of voxels keeping the same NuFiD index of 6 after the filtering. Indeed, for fODF NuFiD indices of 0, 2 and 4, the distribution is always at its maximum where the a-fODF NuFiD index is equal to its initial index. In this case, the maximum of the distribution shifts to 4, and the curve shows that the a-fODF NuFiD indices are mostly distributed between 2 and 5.

This decrease is even more striking for fODF NuFiD indices of 8 (purple curve) and 10 (brown curve) where we report close to no voxels keeping their original NuFiD index in the a-fODF image. This means that for fODFs with 6 or more fiber directions, the neighbourhood rarely supports as many directions.

The curves also highlight that the filtering has a tendency to attenuate the principal fiber directions rather than to extract new fiber directions. However, for voxels with an initial NuFiD index between 0 and 4, there is always a small proportion of voxels increasing their NuFiD index by 1.

### 4.3. Template and proportions of asymmetries

The template of asymmetries, obtained by averaging all asymmetry maps registered to MNI space, is shown in Figure 13 for sagittal and coronal views. As expected, ASI is at its highest near the cortex, where a-fODFs bend, branch and display lobes of uneven probability (endpoint a-fODFs). The other regions of asymmetries identified in Figure 9 are also visible on the template. The voxels located at the exit of the CC, slightly above and below the core of the bundle where projection and commissural fiber pathways intersect (red arrow on the figure), consistently display a high ASI on the whole extent of the ventricles along the coronal axis. The core of major, “easy-to-track” fiber bundles, like the corticospinal tract (CST) and the CC, do not display a high ASI. This is expected, as the fODFs found at the core of the bundles have a very organized neighbourhood.

**Figure 13:**
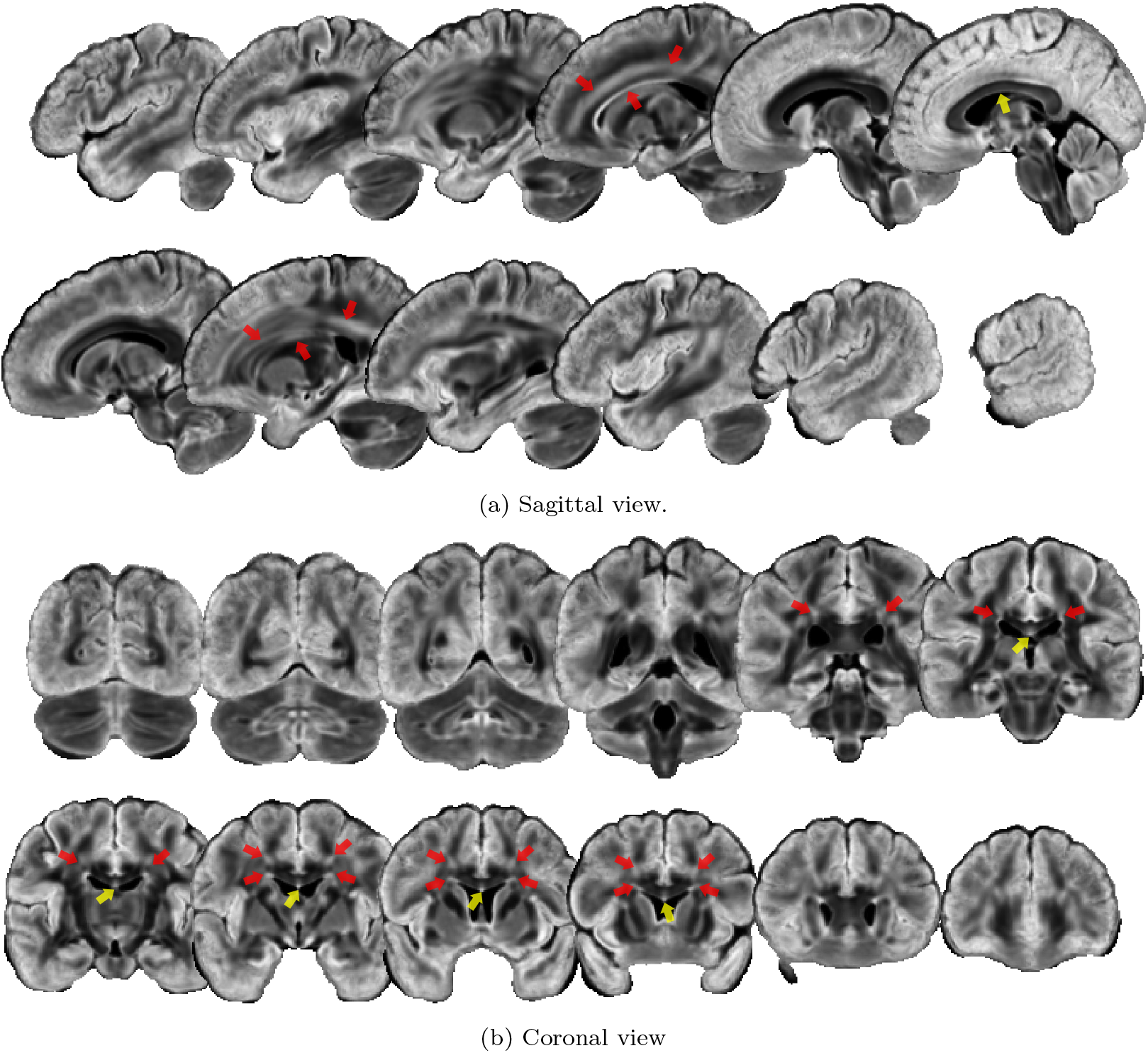
Brain-masked ASI template by slice increments of 10. The colormap goes from black (0, symmetric) to white (0.7879, highly asymmetric). The cortex displays a high degree of asymmetry, as do the bottom-most part of the bridge of the CC (yellow arrow) and the region slightly above and below the exit of the bridge of the CC (red arrow).

Figure 14 shows the distribution of voxel-wise standard deviations inside the WM mask (14a), inside the GM (14b) and for the union of the GM and WM masks (14c). First of all, we note that the std inside the WM mask is low, with very few voxels having a std higher than 0.15. On the contrary, we report a much bigger variability inside voxels classified as GM. There are still a lot of voxels with std < 0.15, and when putting together WM and GM voxels, we see that there is a majority of voxels where std < 0.15. The voxel-wise standard deviation is shown in sub-figure 14d using a grayscale colormap. We see that the std is indeed at its highest along the cortex, and that it progressively goes down as we reach deeper into the WM. We also note some variability around the ventricles.

**Figure 14:**
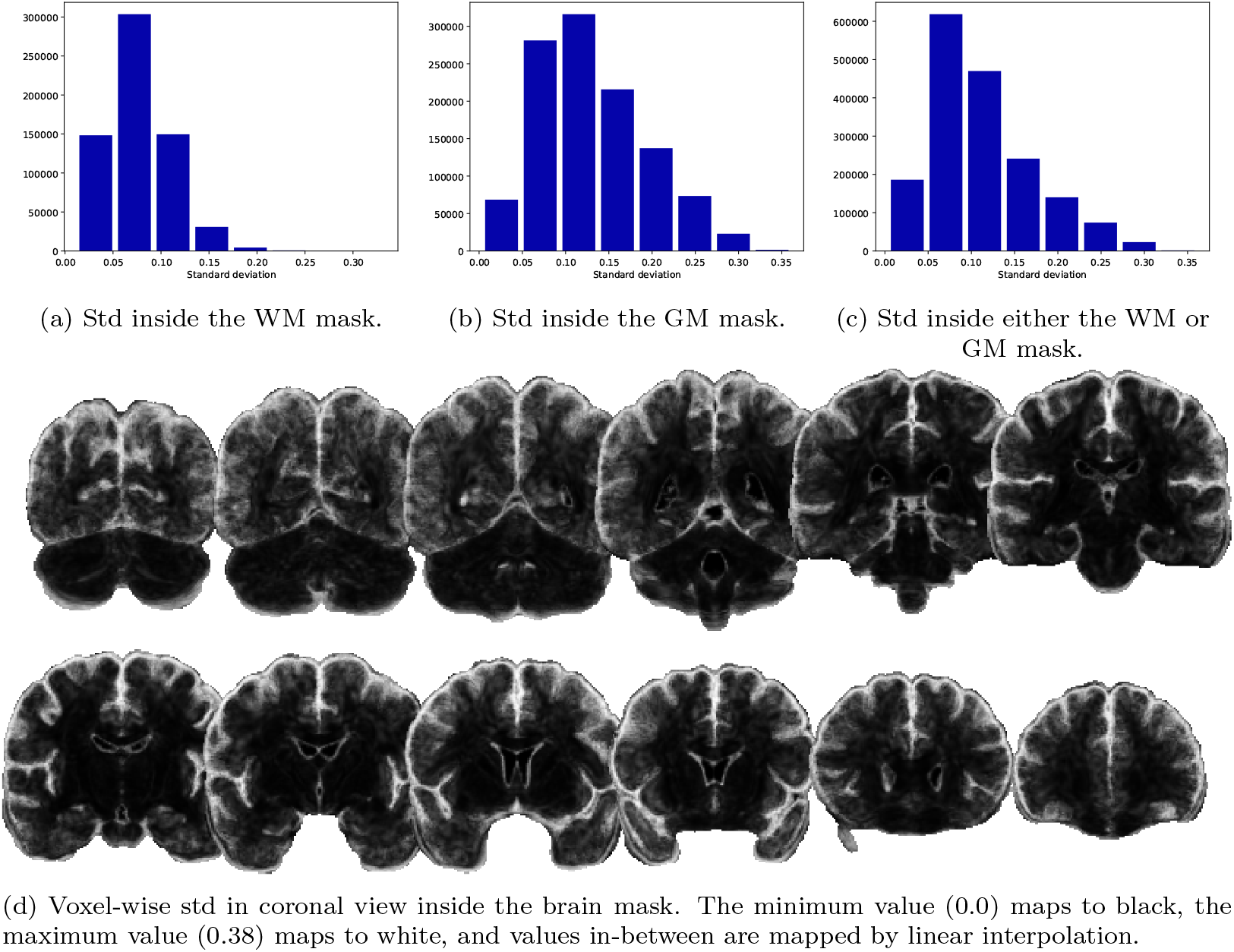
Distribution of std of voxel-wise asymmetry degree across subjects. A low std means less variability across subjects.

In Figure 15 we report the percentage of voxels with an ASI higher than a varying threshold in the range [0.0, 0.8]. The blue curve considers only voxels classified as WM and the yellow one, GM voxels. The green curve includes both WM and GM, while the red one includes all brain voxels as defined by the MNI brain mask (which includes the CSF and ventricles). The curve for the union of WM and GM is well aligned to the one for the whole brain, meaning that the voxels corresponding to ventricles/CSF were correctly identified as such during MSMT-CSD fODF reconstruction, and hence clipped to 0 post-filtering. It is expected that the green curve would be slightly above the orange one, as the brain mask contains more voxels than the union of the GM and WM masks. By defining the set of voxels with an asymmetry degree higher than the threshold as *asymmetric voxels*, we report higher proportions of asymmetric voxels among GM voxels than among WM voxels. This is consistent with the values reported in Figure 13, where the asymmetry degree is higher near the cortex than in the deep WM. By fixing to 0.35 the asymmetry degree threshold, we report 43.49% of asymmetric voxels inside the WM mask, 73.25% inside the GM mask and 62.43% inside either GM or WM.

**Figure 15:**
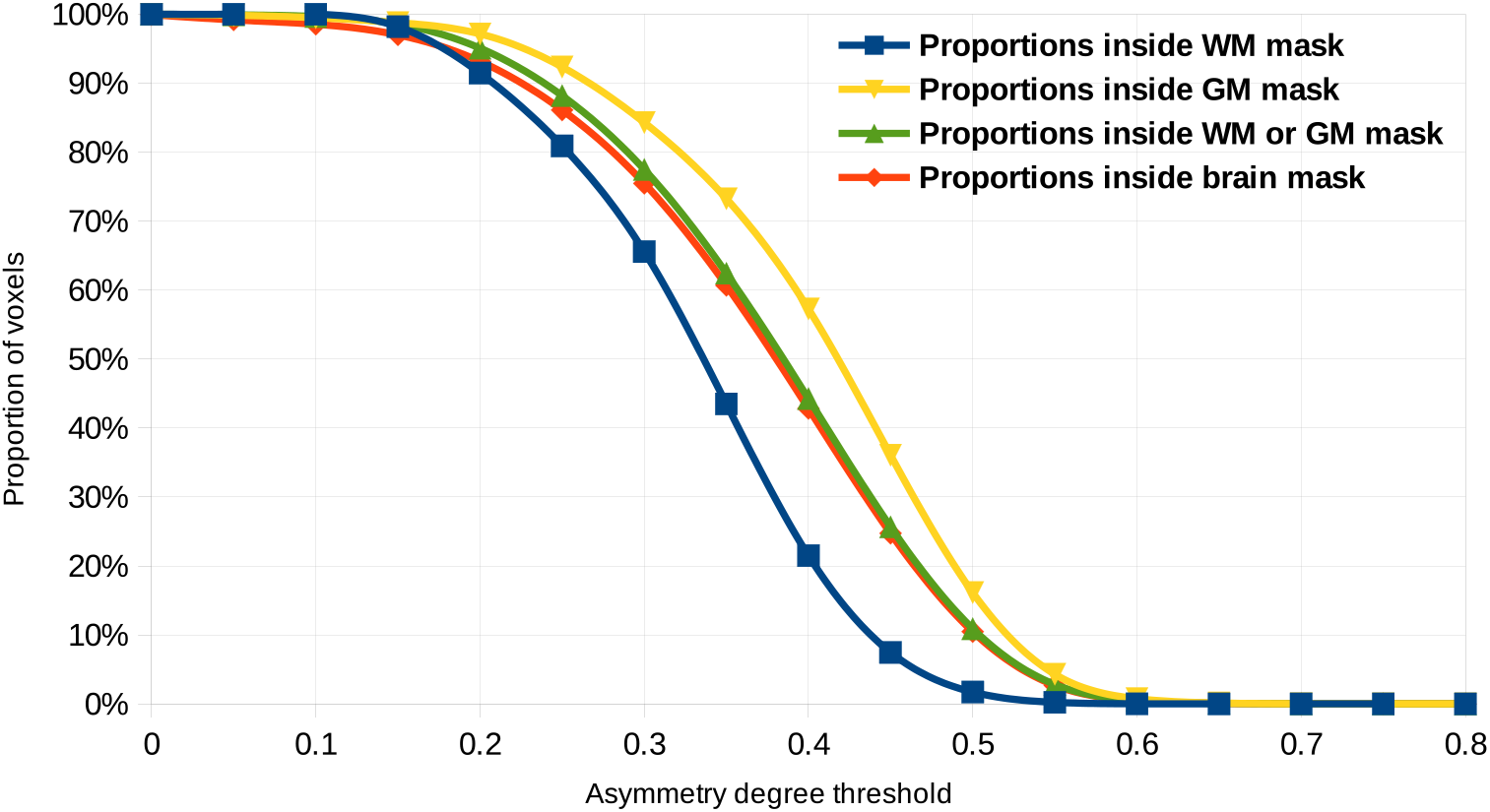
Proportions of voxels (in percents) with an asymmetry degree greater than the threshold (*x* axis) inside the WM (blue), GM (yellow), either WM or GM (green) and inside the brain mask (orange).

In Figure 16, voxels classified as asymmetric are rendered on top of an anatomical T1-weighted image. Asymmetric WM voxels are drawn in yellow and asymmetric GM voxels are drawn in green. This result confirms that the chosen threshold correctly labels the interesting regions of asymmetries previously reported as such. These regions include the periventricular area located above and below the exit of the CC bridge, the bottom-most portion of the CC bridge, the gyral crowns and the cortex.

**Figure 16:**
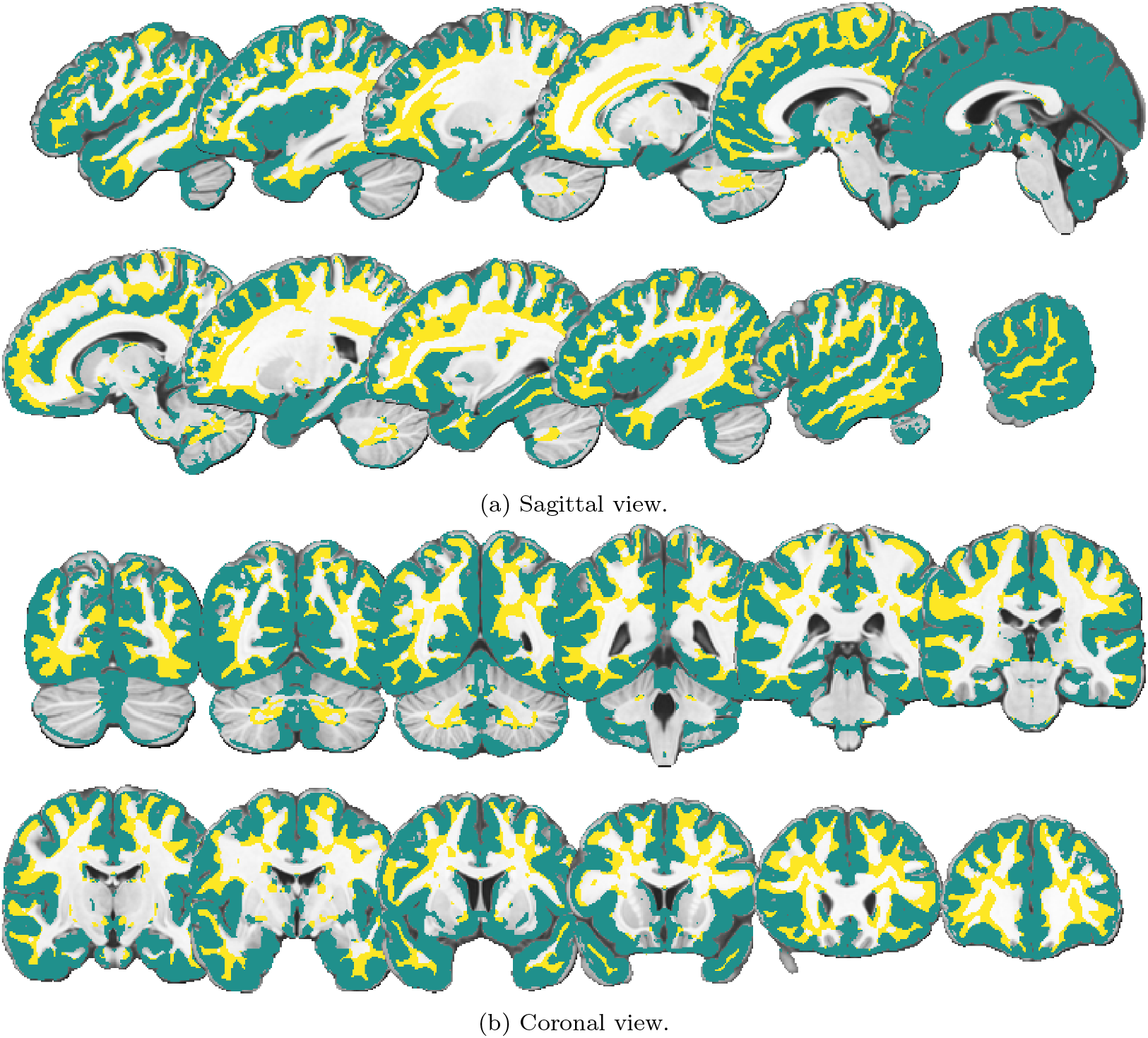
Asymmetric regions (ASI > 0.35) are displayed on top of a T1-weighted image. Asymmetric voxels classified as WM are drawn in yellow and those classified as GM are drawn in green.

On a desktop computer with an Intel Core i9-9900K CPU and a NVIDIA GeForce GTX1660 GPU, the filtering completes in less than 1.5 minutes on a whole brain with isotropic voxels of 1.25mm^3^ using the parameters described above. In comparison, a single-threaded, pure-python implementation requires around 2 hours to complete for the same data and parameters. When enabling angle filtering, and therefore the summation on sphere directions *v*, our GPU-accelerated implementation requires nearly 25 minutes to complete.

## 5. Discussion

In this work, we proposed a novel unified equation for estimating asymmetric ODFs from an input symmetric ODF field. Our edge-preserving, non-iterative filter successfully estimates asymmetric configurations, showing greater inter-voxel coherence than symmetric ODFs, in a few minutes thanks to our efficient GPU-accelerated implementation. We also built a new template from 21 test/retest MSMT-CSD fODF reconstructions from the HCP database describing the voxel-wise ASI. We consistently identified organized, asymmetric regions inside the brain, which account for 43.49% of WM voxels and 73.25% of GM voxels. We also proposed a new measure describing the number of fiber directions, showing new contrasts in a-ODF images. Because the lack of an easy-to-use, fast and freely available implementation of asymmetric filtering algorithms is a major drawback for the wider adoption of asymmetric ODFs, our method is distributed as a minimal, open-source application. Our GPU-accelerated implementation runs in less than 1.5 minutes on a full brain with voxel size of 1.25 mm^3^.

### 5.1. Filter design and selection of hyperparameters

Our filtering method relies on Gaussian weights for all regularization terms. This well-understood function is a straightforward choice in image processing, as it results in smooth transitions between voxels, and gives a greater importance to voxels close to the current voxel *x* — be it in terms of spatial distance, angular distance or difference in intensities — than to farther ones. As we recall, Barmpoutis et al [8] used von Mises-Fisher distributions for weighting by the angle between sampled directions and the alignment between a sampled direction and the direction to a neighbour. However, since both the von Mises-Fisher and Gaussian distributions follow the formulation exp (*f*(*x*)), a similar weighting could be obtained from both distributions by adequately tuning hyperparameter values. While it is not obvious from Figure 5 that the Gaussian weights are useful for spatial regularization, it makes sense in real-life scenarios, where voxels closer to each other have a higher probability of belonging to the same fiber bundle than further ones. Range filtering is also of particular interest for reducing the contribution of the background signal to the a-ODF estimation. As we saw in Figure 6, *R_angle_* smooths SF directions between themselves. Because MSMT-CSD fODFs are already sharp and clean, this behaviour is not useful as it only results in a loss of angular resolution at the cost of considerably higher computation times. The values chosen for each *σ*_{·}_ were chosen based on the results obtained on the test data and validated on a subset of our cohort in order to obtain a slight smoothing effect on the input fODF image.

The use of an *x*-centered window instead of a conic neighbourhood is motivated by its simplicity of implementation and easier generalization. The choice of a conic neighbourhood is over-designed for the specific purpose of generating asymmetric ODFs, and requires defining as many neighbourhoods as there are sampled sphere directions. The use of a window of fixed width mimics more closely the behaviour of a classical Gaussian or bilateral filtering; disabling *R_angle_* and *R_align_* would be equivalent to applying a bilateral filter on each SF direction, which could act as a symmetric ODF denoising method. Furthermore, the size of our neighbourhood is controlled solely by *σ_spatial_* and does not require fixing any additional hyperparameters such as a cone radius and height.

In [14], the authors report competitive results using a cone model regularization inspired by steerable filters. Because there is no clear distinction between spatial, alignment, angle or range regularization in the context of steerable filters, it remains to be seen how this approach would fit into our equation. However, at first glance, they do not report results that we could not reproduce with our method.

### 5.2. Asymmetric fiber configurations

Our method estimates convincing asymmetric fiber configurations such as branching, fanning, bending and endpoint a-fODFs, all of which were also reported by previous authors. Furthermore, we observe behaviours in the gyral blades similar to those reported using a more complex reconstruction method such as [17]. The debate of filtering versus optimization methods for asymmetric ODFs is still an open question, as no study has effectively compared these two approaches yet. Still, our filtering equation does a good job at generalizing the previously proposed methods, and the code availability of our application makes it all the more suited for integrating and studying more filtering variants. It could even be used as the ground work for comparing those two popular approaches.

A majority of voxels having an initial NuFiD index of 6 or more (NuFO of 3 and above) gets downgraded to a lower NuFiD index, suggesting that our filtering removes fiber directions which are not supported by the neighbourhood, being a consequence of the symmetry constraint imposed on the input fODF. Rarely, our filtering method extracts new significant fiber directions from the input symmetric fODF field. The rare occurrence of these events is not surprising, as the proposed method relies on Gaussian functions, which act as low-pass filters and always assign a greater weight to the current voxels than to its neighbours. To overcome this limitation, we would need to design a different filtering kernel relying on more advanced concepts. For instance, SE(3)-based approaches have been proposed for ODF filtering [9, 37, 38, 39, 40, 41], and although none of them has been used for the specific purpose of generating asymmetric ODFs, they offer a promising framework for this task. However, these formulations rely on sophisticated mathematical concepts, which are beyond the scope of the current work.

### 5.3. Interpretation of our ASI template

In [13], the authors show a map quantifying the asymmetry of a-fODFs estimated from solving an optimization problem. They report asymmetric regions which are consistently observed across multiple subjects. Comparing their results with ours, we see that the asymmetric regions reported in both works are in strong agreement. Indeed, the bottom and exit of the bridge of the CC and the gyral blades are described as asymmetric in both works. They use their own measure of asymmetry, which is not normalized, and a smaller cohort of 10 subjects. Hence, our proposed template validates the asymmetric regions reported previously and, once again, suggests that our filtering method behaves similarly to a more complex optimization method.

Because we report a considerable proportion of asymmetric voxels (62.43% of WM and GM) following a slight, edge-preserving averaging of our input symmetric fODF images, being aware of their happening is of major interest; it is not an exceptional event of low occurrence and it is likely to impact the reconstruction of a high proportion of WM pathways. The regions of asymmetry reported in our work could serve as a basis for studying particular WM regions such as those highlighted in figures 13 and 16. As seen in sub-figure 14b, the std inside GM voxels is greater than for WM voxels, suggesting we should be careful about how we interpret results inside GM. However, we hypothesize that this variability is mostly a consequence of registration errors introducing noise at the interface between GM and CSF. This is expected when fitting the cortex of each subject to a standard template. The same can be said for the high std around the ventricles. The strength of MSMT-CSD is its ability to resolve more precise fODF at tissue interfaces. As such, and as can be seen in Figure 9, the asymmetries reported in the GM are also of interest despite the reported std.

As a limitation, we note that the ASI template is not perfectly aligned to the MNI T1 image, being slightly squeezed along the Z-axis (this is particularly noticeable in Figure 16). The result could be improved by registering T1-weighted images in their native resolution of 0.7 mm^3^ directly to the MNI template, instead of using the 1.25 mm^3^ resampled ones. Indeed, the finer details of the original acquisitions could prove helpful when registering the cortex and ventricles. This could potentially reduce the standard deviations reported for these regions. However, because it is not compared to other modalities aligned to the MNI template, we hypothesize that it is of no consequences for the current study. Still, it is worth noting that, in its current state, our template should be used carefully with other MNI-registered images.

### 5.4. Future works

We are optimistic a-fODFs obtained with our method could help tracking at the WM/GM interface, as it has been shown in [17]. Also, because a lower number of possible fiber directions gives a better idea of the underlying anatomy, a-fODFs offer great promises for improving the reconstruction of fiber pathways. A-fODF-based tractography [10, 11, 13, 17] is still an open problem, as there is no consensus yet on how a good a-fODF-based tracking algorithm should behave. Now that there is a novel, open-source framework for asymmetric ODF estimation, we hope to tackle the problem of tracking on these asymmetric spherical functions. Also, we wish to develop a more data-driven approach for fixing the optimal hyperparameter values for our filter weights. In particular, the optimal hyperparameter values might vary based on the content of the region where the filtering is applied. As it has been pleaded in [10, 11], asymmetric ODFs offer great opportunities for enhancing signals of low resolution, which is typically the case in clinical settings. It would therefore be interesting to study the effect of our filtering algorithm on these types of acquisitions, and even on other signal representations such as the diffusion ODF [42, 2]. A comparative study of asymmetric ODF reconstruction versus filtering methods would also be interesting. These could even be compared to more advanced representations of the fiber distribution, such as fiber trajectory distributions (FTD) [16].

## 6. Conclusion

In this work, we proposed a new open-source, GPU-accelerated python application for estimating asymmetric ODFs from an input symmetric ODF image. Using our novel unified equation together with measures for describing asymmetric ODFs, such as the asymmetry index (ASI) and our novel number of fiber directions (NuFiD), we described *where* and *how* asymmetric ODF configurations occur inside diffusion-weighted brain acquisitions. Now that it has been showed that asymmetric ODFs happen in more than 60% of gray and white matter voxels, we hope that our easy-to-use “plug-and-play” application will encourage the diffusion MRI community to go beyond the symmetric ODF representation.

## 7. Acknowledgements

The authors thank Philippe Karan, from the Sherbrooke Connectivity Imaging Laboratory, for the multi-shell multi-tissue fiber ODF reconstructions used in this work.

The authors also thank the National Science and Engineering Research Council of Canada (NSERC) for funding this research, through the Discovery Grants (DG) program and the Canada Graduate Scholarships - Master’s (CSG M) program. Fundings were also provided in part by the Research Chair in Neuroscience and by the Excellence Scholarship program from the Université de Sherbrooke.

Data were provided in part by the Human Connectome Project, WU-Minn Consortium (Principal Investigators: David Van Essen and Kamil Ugurbil; 1U54MH091657) funded by the 16 NIH Institutes and Centers that support the NIH Blueprint for Neuroscience Research; and by the McDonnell Center for Systems Neuroscience at Washington University.

## References

[1] P. J. Basser, J. Mattiello, D. Lebihan, Estimation of the Effective Self-Diffusion Tensor from the NMR Spin Echo, Journal of Magnetic Resonance, Series B 103 (3) (1994) 247–254. doi: 10.1006/jmrb.1994.1037.

[2] M. Descoteaux, High Angular Resolution Diffusion MRI: from Local Estimation to Segmentation and Tractography, Ph.D. thesis, Université Nice-Sophia Antipolis (Feb. 2008).

[3] J. D. Tournier, F. Calamante, D. G. Gadian, A. Connelly, Direct estimation of the fiber orientation density function from diffusion-weighted MRI data using spherical deconvolution, NeuroImage 23 (3) (2004) 1176–1185, number: 3. doi: 10.1016/j.neuroimage.2004.07.037.

[4] B. Jeurissen, A. Leemans, J.-D. Tournier, D. K. Jones, J. Sijbers, Investigating the prevalence of complex fiber configurations in white matter tissue with diffusion magnetic resonance imaging, Human Brain Mapping 34 (11) (2013) 2747–2766. doi: 10.1002/hbm.22099.

[5] S. Jbabdi, H. Johansen-Berg, Tractography: Where Do We Go from Here?, Brain Connect 1 (3) (2011) 169–183. doi: 10.1089/brain.2011.0033.

[6] K. H. Maier-Hein, P. F. Neher, J.-C. Houde, M.-A. Côté, E. Garyfallidis, J. Zhong, M. Chamberland, F.-C. Yeh, Y.-C. Lin, Q. Ji, W. E. Reddick, J. O. Glass, D. Q. Chen, Y. Feng, C. Gao, Y. Wu, J. Ma, R. He, Q. Li, C.-F. Westin, S. Deslauriers-Gauthier, J. O. O. González, M. Paquette, S. St-Jean, G. Girard, F. Rheault, J. Sidhu, C. M. W. Tax, F. Guo, H. Y. Mesri, S. Dávid, M. Froeling, A. M. Heemskerk, A. Leemans, A. Boré, B. Pinsard, C. Bedetti, M. Desrosiers, S. Brambati, J. Doyon, A. Sarica, R. Vasta, A. Cerasa, A. Quattrone, J. Yeatman, A. R. Khan, W. Hodges, S. Alexander, D. Romascano, M. Barakovic, A. Auría, O. Esteban, A. Lemkaddem, J.-P. Thiran, H. E. Cetingul, B. L. Odry, B. Mailhe, M. S. Nadar, F. Pizzagalli, G. Prasad, J. E. Villalon-Reina, J. Galvis, P. M. Thompson, F. D. S. Requejo, P. L. Laguna, L. M. Lacerda, R. Barrett, F. Dell’Acqua, M. Catani, L. Petit, E. Caruyer, A. Daducci, T. B. Dyrby, T. Holland-Letz, C. C. Hilgetag, B. Stieltjes, M. Descoteaux, The challenge of mapping the human connectome based on diffusion tractography, Nature Communications 8 (1) (2017) 1349. doi: 10.1038/s41467-017-01285-x.

[7] S. Delputte, H. Dierckx, E. Fieremans, Y. D’Asseler, R. Achten, I. Lemahieu, Postprocessing of brain white matter fiber orientation distribution functions, in: 2007 4th IEEE International Symposium on Biomedical Imaging: From Nano to Macro, 2007, pp. 784–787, iSSN: 1945-8452. doi: 10.1109/ISBI.2007.356969.

[8] A. Barmpoutis, B. C. Vemuri, D. Howland, J. R. Forder, Extracting tractosemas from a displacement probability field for tractography in dw-mri, in: D. Metaxas, L. Axel, G. Fichtinger, G. Székely (Eds.), Medical Image Computing and Computer-Assisted Intervention – MICCAI 2008, Lecture Notes in Computer Science, Springer, Berlin, Heidelberg, 2008, pp. 9–16. doi: 10.1007/978-3-540-85988-8\_2.

[9] R. Duits, E. Franken, Left-Invariant Diffusions on the Space of Positions and Orientations and their Application to Crossing-Preserving Smoothing of HARDI images, International Journal of Computer Vision 92 (3) (2011) 231–264. doi: 10.1007/s11263-010-0332-z.

[10] H.-H. Ehricke, K.-M. Otto, U. Klose, Regularization of bending and crossing white matter fibers in MRI Q-ball fields, Magnetic Resonance Imaging 29 (7) (2011) 916–926. doi: 10.1016/j.mri.2011.05.002.

[11] M. Reisert, E. Kellner, V. G. Kiselev, About the geometry of asymmetric fiber orientation distributions, IEEE Transactions on Medical Imaging 31 (6) (2012) 1240–1249. doi: 10.1109/TMI.2012.2187916.

[12] J. S. W. Campbell, P. MomayyezSiahkal, P. Savadjiev, I. R. Leppert, K. Siddiqi, G. B. Pike, Beyond Crossing Fibers: Bootstrap Probabilistic Tractography Using Complex Subvoxel Fiber Geometries, Front. Neurol. 5, publisher: Frontiers (2014). doi: 10.3389/fneur.2014.00216.

[13] M. Bastiani, M. Cottaar, K. Dikranian, A. Ghosh, H. Zhang, D. C. Alexander, T. E. Behrens, S. Jbabdi, S. N. Sotiropoulos, Improved tractography using asymmetric fibre orientation distributions, NeuroImage 158 (2017) 205–218. doi: 10.1016/j.neuroimage.2017.06.050.

[14] S. Cetin Karayumak, E. Ozarslan, G. Unal, Asymmetric Orientation Distribution Functions (AODFs) revealing intravoxel geometry in diffusion MRI, Magnetic Resonance Imaging 49 (2018) 145–158. doi: 10.1016/j.mri.2018.03.006.

[15] Y. Wu, W. Lin, D. Shen, P.-T. Yap, Asymmetry Spectrum Imaging for Baby Diffusion Tractography, in: A. C. S. Chung, J. C. Gee, P. A. Yushkevich, S. Bao (Eds.), Information Processing in Medical Imaging, Lecture Notes in Computer Science, Springer International Publishing, Cham, 2019, pp. 319–331. doi: 10.1007/978-3-030-20351-1\_24.

[16] Y. Feng, J. He, Asymmetric fiber trajectory distribution estimation using streamline differential equation, Medical Image Analysis 63 (2020) 101686. doi: 10.1016/j.media.2020.101686.

[17] Y. Wu, Y. Hong, Y. Feng, D. Shen, P.-T. Yap, Mitigating gyral bias in cortical tractography via asymmetric fiber orientation distributions, Medical Image Analysis 59 (2020) 101543. doi: 10.1016/j.media.2019.101543.

[18] B. Jeurissen, J.-D. Tournier, T. Dhollander, A. Connelly, J. Sijbers, Multi-tissue constrained spherical deconvolution for improved analysis of multi-shell diffusion MRI data, NeuroImage 103 (2014) 411–426. doi: 10.1016/j.neuroimage.2014.07.061.

[19] F. Dell’Acqua, A. Simmons, S. C. R. Williams, M. Catani, Can spherical deconvolution provide more information than fiber orientations? Hindrance modulated orientational anisotropy, a true-tract specific index to characterize white matter diffusion, Human Brain Mapping 34 (10) (2013) 2464–2483. doi: 10.1002/hbm.22080.

[20] C. Tomasi, R. Manduchi, Bilateral filtering for gray and color images, in: Sixth International Conference on Computer Vision (IEEE Cat. No.98CH36271), 1998, pp. 839–846. doi: 10.1109/ICCV.1998.710815.

[21] M. Petrou, C. Petrou, Image Processing: The Fundamentals, 2nd Edition, John Wiley & Sons, Ltd, 2010. doi: 10.1002/9781119994398. URL https://onlinelibrary.wiley.com/doi/10.1002/9781119994398

[22] A. Klöckner, N. Pinto, Y. Lee, B. Catanzaro, P. Ivanov, A. Fasih, PyCUDA and PyOpenCL: A scripting-based approach to GPU runtime code generation, Parallel Computing 38 (3) (2012) 157–174. doi: 10.1016/j.parco.2011.09.001.

[23] S. K. Lam, A. Pitrou, S. Seibert, Numba: a LLVM-based Python JIT compiler, in: Proceedings of the Second Workshop on the LLVM Compiler Infrastructure in HPC, LLVM ’15, Association for Computing Machinery, New York, NY, USA, 2015, pp. 1–6. doi: 10.1145/2833157.2833162.

[24] J.-D. Tournier, F. Calamante, A. Connelly, Robust determination of the fibre orientation distribution in diffusion MRI: Non-negativity constrained super-resolved spherical deconvolution, NeuroImage 35 (4) (2007) 1459–1472, number: 4. doi: 10.1016/j.neuroimage.2007.02.016.

[25] P. Fillard, M. Descoteaux, A. Goh, S. Gouttard, B. Jeurissen, J. Malcolm, A. Ramirez-Manzanares, M. Reisert, K. Sakaie, F. Tensaouti, T. Yo, J.-F. Mangin, C. Poupon, Quantitative evaluation of 10 tractography algorithms on a realistic diffusion MR phantom, NeuroImage 56 (1) (2011) 220–234. doi: 10.1016/j.neuroimage.2011.01.032.

[26] C. Poupon, L. Laribiere, G. Tournier, J. Bernard, D. Fournier, P. Fillard, M. Descoteaux, J.-F. Mangin, A Diffusion Hardware Phantom Looking Like a Coronal Brain Slice, in: ISMRM 18th Scientific Meeting and Exhibition, Stockholm, Sweden, 2010.

[27] C. Poupon, B. Rieul, I. Kezele, M. Perrin, F. Poupon, J.-F. Mangin, New diffusion phantoms dedicated to the study and validation of high-angular-resolution diffusion imaging (HARDI) models, Magnetic Resonance in Medicine 60 (6) (2008) 1276–1283. doi: 10.1002/mrm.21789.

[28] P. F. Neher, M.-A. Côté, J.-C. Houde, M. Descoteaux, K. H. Maier-Hein, Fiber tractography using machine learning, NeuroImage 158 (2017) 417–429. doi: 10.1016/j.neuroimage.2017.07.028.

[29] D. C. Van Essen, S. M. Smith, D. M. Barch, T. E. J. Behrens, E. Yacoub, K. Ugurbil, The WU-Minn Human Connectome Project: An overview, NeuroImage 80 (2013) 62–79. doi: 10.1016/j.neuroimage.2013.05.041.

[30] E. Garyfallidis, M. Brett, B. Amirbekian, A. Rokem, S. Van Der Walt, M. Descoteaux, I. Nimmo-Smith, Dipy, a library for the analysis of diffusion MRI data, Frontiers in Neuroinformatics 8 (2014) 8. doi: 10.3389/fninf.2014.00008.

[31] M. F. Glasser, S. N. Sotiropoulos, J. A. Wilson, T. S. Coalson, B. Fischl, J. L. Andersson, J. Xu, S. Jbabdi, M. Webster, J. R. Polimeni, D. C. Van Essen, M. Jenkinson, The minimal preprocessing pipelines for the Human Connectome Project, NeuroImage 80 (2013) 105–124. doi: 10.1016/j.neuroimage.2013.04.127.

[32] G. Theaud, J.-C. Houde, A. Boré, F. Rheault, F. Morency, M. Descoteaux, TractoFlow: A robust, efficient and reproducible diffusion MRI pipeline leveraging Nextflow & Singularity, NeuroImage 218 (2020) 116889. doi: 10.1016/j.neuroimage.2020.116889.

[33] C. Poirier, M. Descoteaux, G. Gilet, Accelerating Geometry-Based Spherical Harmonics Glyphs Rendering for dMRI Using Modern OpenGL, in: S. Cetin-Karayumak, D. Christiaens, M. Figini, P. Guevara, N. Gyori, V. Nath, T. Pieciak (Eds.), Computational Diffusion MRI, Lecture Notes in Computer Science, Springer International Publishing, Cham, 2021, pp. 144–155. doi: 10.1007/978-3-030-87615-9\_13.

[34] V. Fonov, A. Evans, R. McKinstry, C. Almli, D. Collins, Unbiased nonlinear average age-appropriate brain templates from birth to adulthood, NeuroImage 47 (2009) S102. doi: 10.1016/S1053-8119(09)70884-5.

[35] V. Fonov, A. C. Evans, K. Botteron, C. R. Almli, R. C. McKinstry, D. L. Collins, Unbiased average age-appropriate atlases for pediatric studies, NeuroImage 54 (1) (2011) 313–327. doi: 10.1016/j.neuroimage.2010.07.033.

[36] F. Rheault, J.-C. Houde, N. Goyette, F. Morency, M. Descoteaux, Mi-brain, a software to handle tractograms and perform interactive virtual dissection, 2016.

[37] M. Reisert, H. Skibbe, Left-Invariant Diffusion on the Motion Group in terms of the Irreducible Representations of SO(3), arXiv:1202.5414 [cs, math] (Feb. 2012). doi: 10.48550/arXiv.1202.5414. URL http://arxiv.org/abs/1202.5414

[38] R. Duits, T. Dela Haije, E. Creusen, A. Ghosh, Morphological and Linear Scale Spaces for Fiber Enhancement in DW-MRI, J Math Imaging Vis 46 (3) (2013) 326–368. doi: 10.1007/s10851-012-0387-2.

[39] T. C. J. Dela Haije, R. Duits, C. M. W. Tax, Sharpening Fibers in Diffusion Weighted MRI via Erosion, in: C.-F. Westin, A. Vilanova, B. Burgeth (Eds.), Visualization and Processing of Tensors and Higher Order Descriptors for Multi-Valued Data, Mathematics and Visualization, Springer, Berlin, Heidelberg, 2014, pp. 97–126. doi: 10.1007/978-3-642-54301-2_5.

[40] V. Prčkovska, M. Andorrà, P. Villoslada, E. Martinez-Heras, R. Duits, D. Fortin, P. Rodrigues, M. Descoteaux, Contextual Diffusion Image Post-processing Aids Clinical Applications, in: I. Hotz, T. Schultz (Eds.), Visualization and Processing of Higher Order Descriptors for Multi-Valued Data, Mathematics and Visualization, Springer International Publishing, Cham, 2015, pp. 353–377. doi: 10.1007/978-3-319-15090-1_18.

[41] J. M. Portegies, R. H. J. Fick, G. R. Sanguinetti, S. P. L. Meesters, G. Girard, R. Duits, Improving Fiber Alignment in HARDI by Combining Contextual PDE Flow with Constrained Spherical Deconvolution, PLOS ONE 10 (10) (2015) e0138122, publisher: Public Library of Science. doi: 10.1371/journal.pone.0138122.

[42] D. S. Tuch, Q-ball imaging, Magnetic Resonance in Medicine 52 (6) (2004) 1358–1372. doi: 10.1002/mrm.20279.

